# Mapping the Architecture of Protein Complexes in *Arabidopsis* Using Cross-Linking Mass Spectrometry

**DOI:** 10.1101/2025.04.28.651104

**Authors:** Cao Son Trinh, Ruben Shrestha, Pengzhi Mao, William C Conner, Andres V Reyes, Sumudu S Karunadasa, Annie Yu, Grace Liu, Ken Hu, Shou-Ling Xu

## Abstract

Capturing molecular machines in action is essential for understanding protein complex architecture, cellular regulation, and gene function. Here, we present a large-scale structural proteomics resource for *Arabidopsis thaliana* generated using an optimized cross-linking mass spectrometry (XL-MS) workflow. Using the trifunctional cross-linker PhoX, whose phosphonic acid moiety enables immobilized metal affinity chromatography (IMAC)-based enrichment, we selectively enriched cross-linked peptides from whole-cell lysates, chloroplasts, and nuclei. Analysis with pLink 3.2 identified 52,944 unique cross-linked peptide pairs, corresponding to 37,531 residue-level contacts across 5,064 proteins. These data define 3,083 protein–protein interactions, including 2,385 heteromeric and 698 homomultimeric interactions. Comparison with the STRING database showed that 676 interactions are supported by STRING scores ≥0.9. Structural mapping to Protein Data Bank (PDB) and AlphaFold models showed that most cross-links were within the expected 35 Å distance constraint. The dataset further enabled the analysis of protein connectivity and complex topology across diverse molecular assemblies, including the Rubisco holoenzyme, chloroplast 70S ribosome, photosystem complexes, and the cytosolic 80S ribosome together with associated biogenesis and regulatory factors. We also identified histone-associated complexes, including interactions involving an O-acyltransferase. By providing residue-level structural constraints for a substantial portion of the *Arabidopsis* proteome, this study provides a resource for exploring plant molecular machines and their spatial organization.

**Significance Statement:** Understanding how proteins interact within living cells is essential to deciphering cellular architecture and function. However, capturing native protein-protein interactions (PPIs) on a global scale has proven technically challenging. Here, we present a proteome-wide cross-linking mass spectrometry (XL-MS) platform that can systematically map direct PPIs in plant cells without requiring transgenic manipulation. This approach identifies thousands of interactions spanning major subcellular compartments and characterizes the *in situ* organization of critical protein assemblies, such as photosystems, ribosomes, and chromatin-associated connectivity. By mapping both established and less-characterized interactions, this work advances our understanding of the plant protein interactome and provides a valuable resource for investigating the structural organization of the plant proteome.

## Introduction

Understanding how molecular machines operate within cells - and capturing them in action - is essential to unraveling the principles of life and gene function. A powerful tool for this purpose is cross-linking mass spectrometry (XL-MS), an emerging high-throughput approach that enables the capture of protein complexes in their native context across diverse organisms (1, 2). Unlike affinity purification mass spectrometry (AP-MS) and proximity labeling mass spectrometry (PL-MS) (3–5), XL-MS does not require antibodies or genetically tagged transgenes. Instead, it relies on chemical cross-linkers that covalently link spatially proximal residues within or between proteins in solution, followed by protease digestion and mass spectrometric identification of the resulting cross-linked peptides. In addition to generating a "parts list" of interacting proteins, XL-MS provides direct evidence of physical interactions and precisely maps contact sites at the amino acid level, providing valuable structural insight into macromolecular assemblies.

Despite its promise, large-scale proteome-wide XL-MS - such as in whole-cell lysates, isolated organelles, or *in vivo* systems - remains challenging due to the inherent low abundance and heterogeneity of cross-linked peptides and the complexity of data analysis (1, 6). In plants, additional barriers further complicate its application, including rigid cell walls and large central vacuoles that dilute cross-linkers, as well as secondary metabolites that can chemically interfere with cross-linking chemistry (7, 8). Recent pioneering studies in Arabidopsis, algae, and soybean have demonstrated the potential of XL-MS in plant systems (7, 9, 10), but its overall implementation remains limited.

To address these challenges, we developed a cross-linking strategy optimized for proteome-wide mapping in complex Arabidopsis samples using the enrichable cross-linker PhoX. PhoX is a phosphonic acid-based reagent that, like BS3 and DSSO, contains NHS esters and primarily targets lysine residues (11). Its phosphonic acid handle allows efficient enrichment by immobilized metal affinity chromatography (IMAC). With a maximum range of 35Å - consisting of a 20 Å Cα-Cα distance from its rigid structure and an additional 15 Å to account for protein flexibility - PhoX is well suited for large-scale structural proteomics (12). To reduce sample complexity and improve cross-linking efficiency, we used subcellular fractionations. To further improve the detection of the low abundance cross-linked peptides, we performed extensive high-pH reversed-phase HPLC fractionation after IMAC enrichment. Together, these measures allowed for comprehensive proteome-wide XL-MS mapping.

We identified 52,944 unique cross-linked peptide pairs (43,780 intra-protein and 9,164 inter-protein) from cell lysates, chloroplasts, and nuclei. These data supported a heteromeric PPI network of 1,429 proteins and 2,385 interactions, plus 698 homomultimeric interactions. Because STRING catalogs only heteromeric PPIs, we compared that subset: 917 heteromeric interactions meet a confidence score ≥ 0.7, including 676 at ≥ 0.9. The remaining interactions either fell below these thresholds or lacked corresponding entries in STRING, suggesting previously less-characterized physical associations. As many experimental PPIs may not yet be indexed in public databases, our dataset provides a residue-level resource for mapping interaction interfaces and prioritizing candidates for further investigation.

Beyond interaction mapping, our XL-MS data provide structural and functional insights into key cellular processes. By characterizing flexible or transient components within the 70S chloroplast ribosome and photosystem complexes, these results complement existing cryo-EM structures. XL-MS also facilitated the mapping of the photosystem, 80S ribosome, and histone interactomes within heterogeneous protein populations, providing spatial constraints for both established and less characterized associations. We identified an O-acyltransferase that interacts with histones, suggesting a potential mechanism of chromatin regulation via post-translational modification. Together, these findings provide experimental support for protein complex architectures and establish a resource for exploring plant molecular machines.

## Results

### Efficient cross-linking workflow for complex protein samples identifies numerous cross-links in Arabidopsis

To gain structural insights into molecular machines within their native cellular context within Arabidopsis cells, we performed XL-MS (see Methods) on fractionated seedling samples, including cell lysates and chloroplasts from light-grown seedlings and nuclei from dark-grown seedlings. Using the PhoX cross-linker (Fig. 1A), we implemented an optimized pipeline for the proteome-wide detection of low-abundance cross-linked peptides across these complex plant samples.

**Figure 1.**
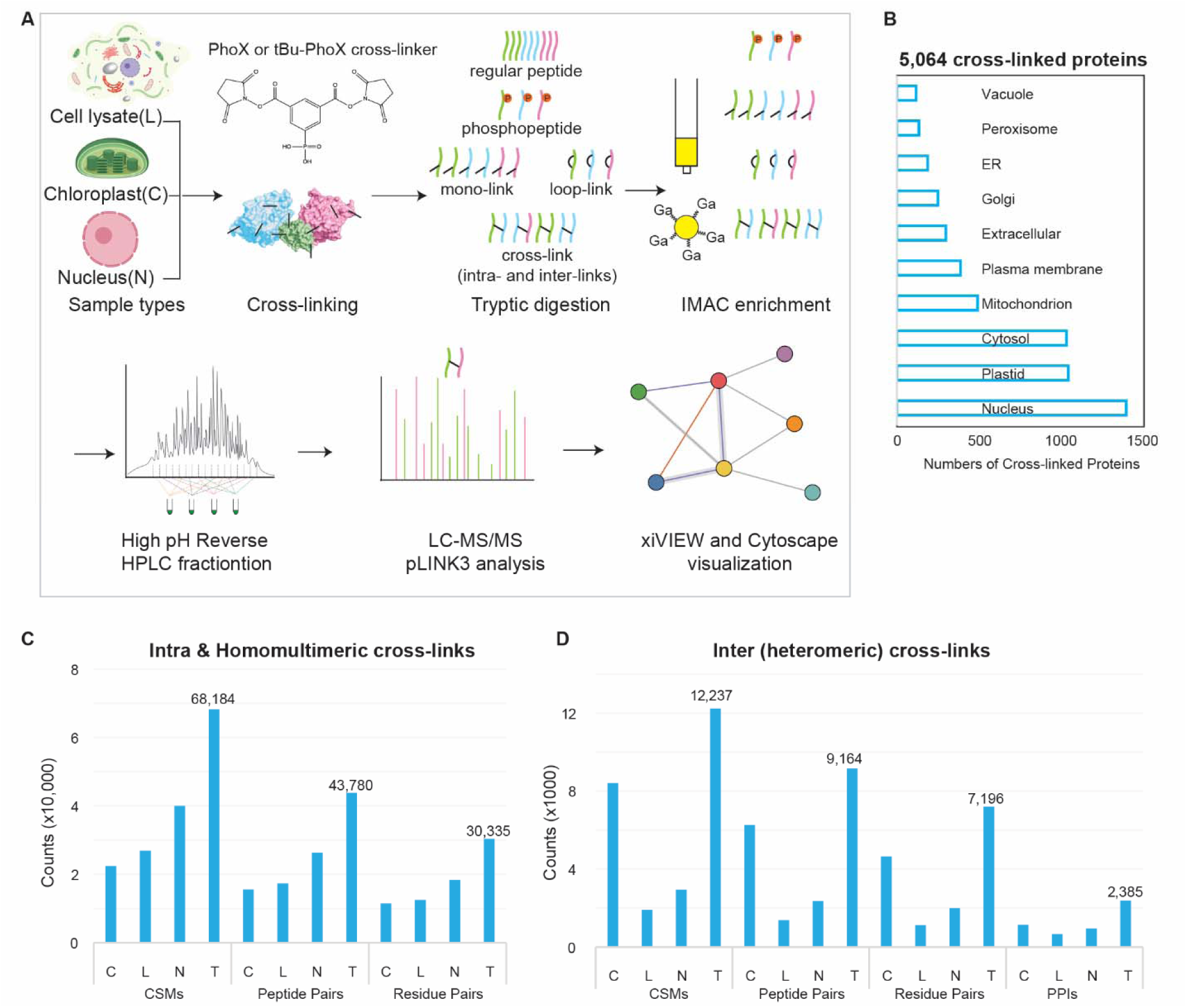
Efficient cross-linking workflow for complex protein samples identifies numerous cross-links in Arabidopsis. (**A**) Schematic of the workflow using PhoX or tBu-PhoX to covalently cross-link spatially proximal lysine residues in Arabidopsis subcellular fractions (lysate, chloroplast, nucleus), followed by digestion, IMAC enrichment, high pH reversed-phase HPLC fractionation, and LC-MS/MS. Data were analyzed using pLINK3.2, Cytoscape, and xiVIEW. (**B**) Subcellular localization of cross-linked proteins based on SUBA5, with redundancy of proteins assigned to multiple compartments. (**C**-**D**) Summary of cross-link identifications. Total and fraction-specific counts for unique intra-protein/homomultimeric (C) and inter-protein/heteromeric (D) cross-links. Data are reported at multiple levels: (C):cross-link spectrum matches (CSMs), peptide pairs, and residue pairs; and (D) the same levels as in (C), plus protein-protein interactions (PPIs). Non-redundant CSMs collapse matches with identical charge,linkage sites, mass, and modifications. FDR (1%) was calculated independently for (C) and (D) at the peptide-pair level.

Optimization of PhoX concentration showed a shift toward higher molecular weights at 2-5 mM (red arrows, Fig. S1A-D), indicating successful cross-linking; as higher concentrations yielded no further improvement, 2 and 5 mM PhoX were selected for subsequent experiments (see Methods). Following cross-linking, proteins were phenol-precipitated, digested with trypsin, and the resulting cross-linked peptides were enriched via GaCl₃-based IMAC (13). Enriched peptides were then fractionated by high-pH reversed-phase chromatography, analyzed by LC-MS/MS on an Orbitrap Eclipse mass spectrometer, utilizing high-resolution MS^1^ and MS^2^ acquisition to ensure reliable cross-link identification. Data were searched using pLink 3.2 (14)(Mao et al., unpublished) and filtered at a 1% peptide-pair FDR. To maximize identification yield, we compared the standard XL workflow with real-time spectral library search (RTLS) (15). While RTLS effectively reduced mono- and loop-link identifications by ∼48% and ∼31%, respectively; it also decreased the detection of our target intra-protein links by 42% and inter-protein links by 40% (Fig. S2A). Consequently, we utilized the standard XL workflow for subsequent chloroplast and nuclear analyses.

Relative abundance analysis confirmed the high enrichment efficiency of the GaCl₃-based IMAC approach (Fig. S2B). Given the potential for FDR propagation from peptide-pair to PPI level identification (16, 17), we further performed searches against entrapment libraries (see Methods) (18). The resulting entrapment-estimated FDP (false discovery proportion) values at the peptide-pair and PPI levels suggested the global error rate remained within acceptable limits (Fig.S2C, Table S1).

We detected 28,677, 21,845, and 18,678 cross-link peptide pairs from the nucleus, chloroplast, and cell lysate, respectively (Fig. 1B-D, S3A, Table S1). In total, we identified 52,944 unique cross-linked peptide pairs (43,780 intra-protein and 9,164 inter-protein) corresponding to 37,531 residue pairs captured across 5,064 proteins and 3,083 protein-protein interactions (2,385 heteromeric and 698 homomultimeric). Subcellular localization analysis via SUBA5 suggests that the cross-linked proteins are distributed across cellular compartments, including the nucleus, plastid, cytosol, mitochondria (Fig. 1B, Table S2). Assessment of protein abundance using PaxDb indicates that our dataset may span a broad dynamic range as cross-links appear to be captured for low- and medium-abundance proteins in addition to highly abundant ones (Fig.S3B).

We observed 51% of heteromeric PPIs were supported by more than one CSM (Fig.S4A). In addition, we observed consistent peptide pairs mapping to the same sites or regions, supporting the reliability of cross-linked peptide identifications (Fig. S4B-F). For example, within ribosomal subunits, we detected site-specific cross-links featuring variable miscleavages, such as those between uL15x-K92 and eL13z-K166/K172 (Fig. S4B). Cross-links with miscleavages were also identified between Histone 1.1-K177/K179 and the SUMO-targeted ubiquitin ligase STUBL1-K187/188 (Fig. S4C). We further observed interactions mapping to conserved sites: the vacuolar proton ATPase subunit VHA-E3-K49 cross-linked to both VHA-B1-K437 and VHA-B2-K438 (Fig.S4D). Similarly, DNA topoisomerase I alpha TOP1A-K672 linked to conserved regions of two dehydrin homologs, ERD14-K38/K42 and COR47-K46 (Fig.S4E), while the 26S proteasome subunit RPN7-K369 linked to the same sites of two homologs, DSS1(I)-K6 and DSS1(V)-K6 (Fig.S4F). Collectively, these observations indicate a high level of agreement within the dataset.

### Validation of intra- and inter-protein cross-links by structural mapping

To assess the validity of the identified cross-links, we mapped intra-protein residue pairs to 736 Arabidopsis PDB structures and 3,575 AlphaFold2 predicted models using ComMap (Table S3) (19). We then evaluated the corresponding Cα-Cα Euclidean distances for these mapped residue pairs. Of the 1,863 and 25,982 successfully mapped cross-links on PDB and AlphaFold structure, respectively, 89.9% (PDB) and 79.5% (AlphaFold) satisfied the 35 Å distance constraint (Fig. 2A–B). As a control, we simulated random lysine residue pairs and mapped them onto both structural datasets. The experimentally mapped cross-link distance distributions differed from the randomized distributions, with a more pronounced difference observed for mappings onto AlphaFold models than for mappings onto PDB structures (Fig. S5A–B) (Fig. S5A-B).

**Figure 2.**
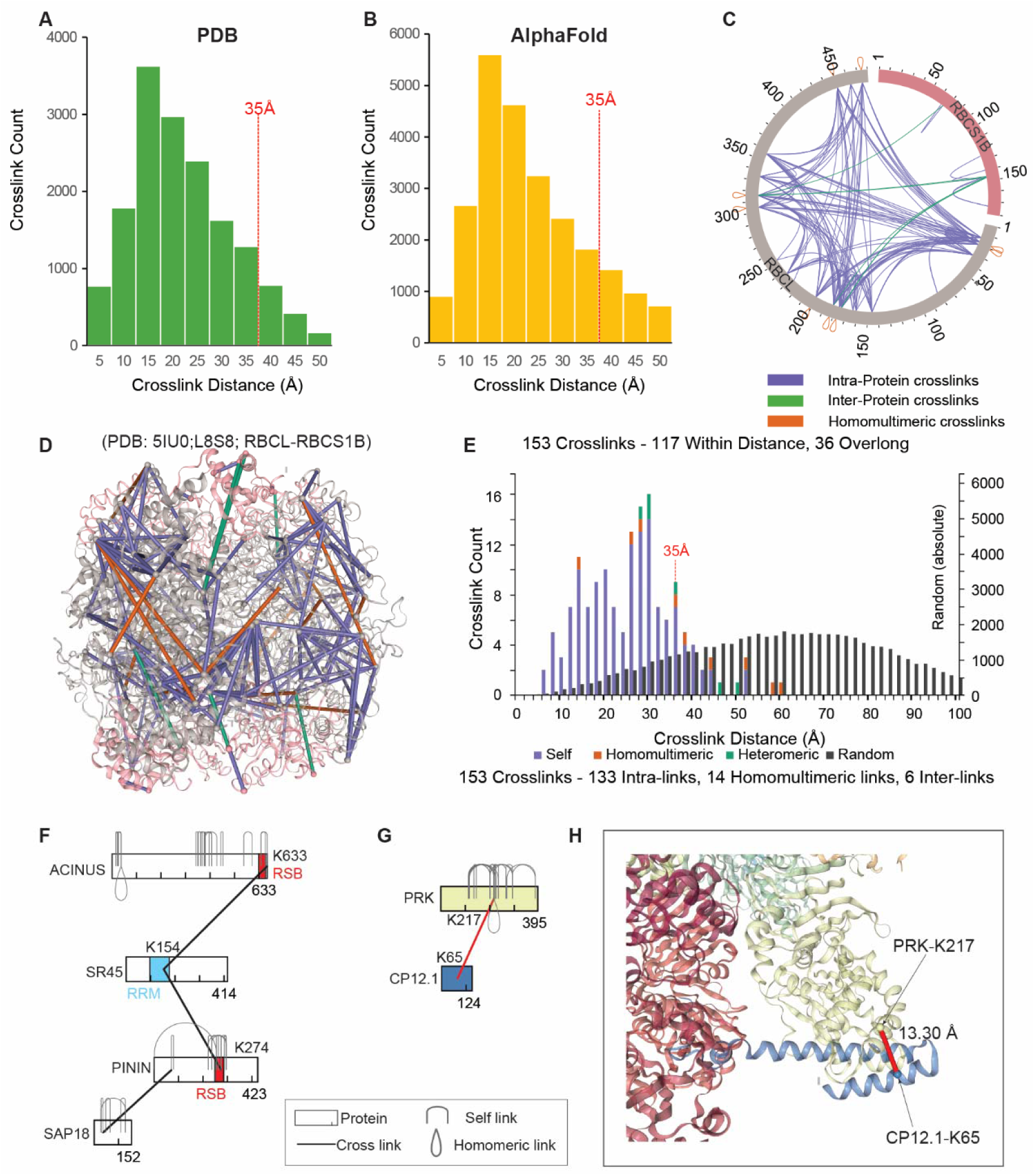
Validation of intra- and inter-protein cross-links by structural mapping. (**A**-**B**) Histogram of Euclidean Cα-Cα distances of intra-protein cross-links mapped to Arabidopsis structures from PDB (A) or AlphaFold (B), showing 96.8% and 94.3% within the 35Å constraint, respectively. (**C**-**E**) Cross-links mapped to the Arabidopsis Rubisco L_8_S_8_ complex modeled from the L_2_S_2_ structure (PDB:5IU0). RBCL is shown in grey and RBCS1B is shown in pink. Self (intra), heteromeric, and homomultimeric links are shown in purple, green, and orange, respectively. The circular view (C) and structural mapping (D-E) show that 76.47% of the cross-links are within 35Å. (**F**) Cross-links between SR45 and ACINUS or PININ occur at known interaction interfaces involving the RRM domain of SR45 and the RSB domain of ACINUS and PININ. ACINUS interacts with SR45 and SAP18 to form the ASAP complex, while PININ interacts with SR45 and SAP18 to form the PSAP complex. (**G-H**) Intra- and inter-protein cross-links between PRK and CP12.1 mapped onto the GAPDH-CP12-PRK complex (PDB:6KEZ), aligning with the known interaction interface.

To further evaluate intra- and inter-protein cross-links, we mapped the detected cross-links onto the Rubisco holoenzyme, revealing conserved connectivity between the RBCL large subunit and multiple RBCS isoforms (Fig.S6A-B). Mapping 153 RBCL–RBCS1B cross-links onto the Arabidopsis Rubisco structure (PDB: 5IU0)(20) showed that 76.4% satisfied the expected ≤ 35 Å Cα-Cα distance constraints, while the remaining cross-links likely reflected structural flexibility (Fig. 2D–E). Similar results were obtained using the RBCL-RBCS1A(PDB:9N37) and RBCL-RBCS1B(PDB:9MUR) structures (Fig. S6A–F).

To assess whether cross-linked lysine residues are positioned within predicted interaction interfaces, we examined several PPIs using available structural data. The cross-link between ACINUS-K633 and SR45-K154 maps near the RSB domain of ACINUS (aa 599-631) and within the RRM domain of SR45 in the ASAP core complex (PDB:4A8X) (21–23) (Fig. 2F, Fig. S7A). Similarly, the cross-link between SR45-K154 and PININ-K274 aligns with the same RRM domain and the predicted RSB domain of PININ (Fig. 2F, Fig. S7B). For PRK-K217 and CP12.1-K65, the cross-link localizes to the interaction interface in the GAPDH-CP12-PRK complex (PDB:6KEZ) (24) with a Cα-Cα distance of 13.3 Å (Fig. 2G-H). These results confirm that the cross-linked residues are located at or within a few amino acids of the interaction interfaces, consistent with previous observations (25).

### XL-MS systematically identifies direct protein-protein interactions and reveals established and less characterized complexes

We imported 3,527 inter-protein cross-links into Cytoscape (26), revealing 3,083 PPIs among 1493 proteins, including 2385 heteromeric and 698 homomultimeric interactions (Fig. 3A, B, Table S1). Proteins were classified using the PANTHER protein class tool, and nodes were colored accordingly (27). Many interacting proteins shared the same classification, supporting the reliability of our PPI map. Notably, 513 of the 1,493 proteins involved in heteromeric interactions lacked PANTHER annotations, contributing to 1,397 interactions - 591 between unclassified proteins and 806 between classified and unclassified proteins.

**Figure 3.**
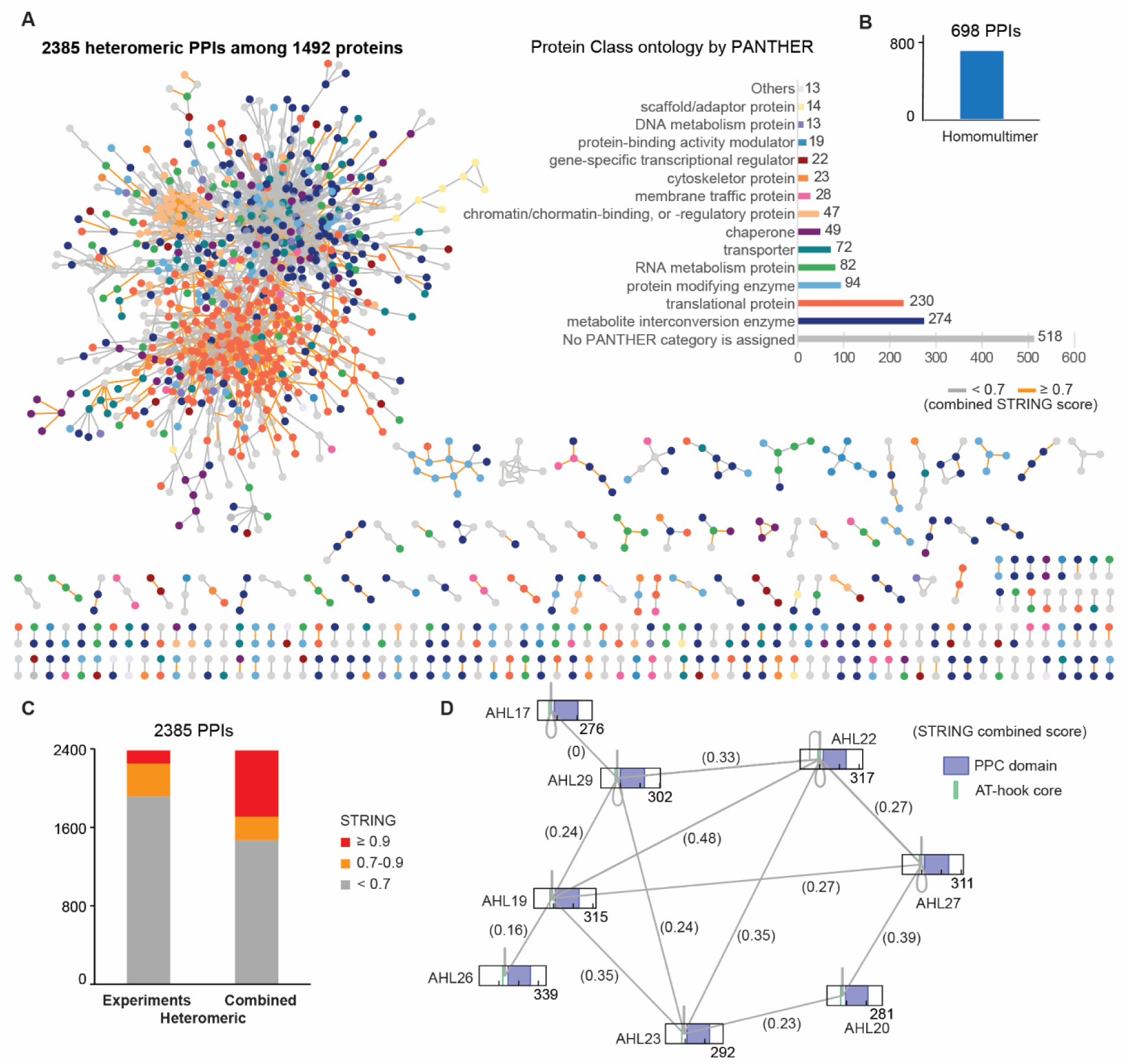
XL-MS identifies direct protein-protein interactions and reveals established and less characterized complexes. (**A-B**) XL-MS identified an Arabidopsis interactome comprising 3,083 PPIs, including 2,385 heteromeric interactions among 1,492 proteins and 698 homomultimeric interactions. Heteromeric interactions were visualized using Cytoscape, with PANTHER protein class annotations mapped onto nodes and STRING scores overlaid on edges. (**C**) Summary of the STRING score, showing combined STRING score and experimental score of all heteromeric PPIs. XL-MS provides experimental support for 80.3% of the heteromeric PPIs that either have lower confidence scores (<0.7) or lack prior experimental evidence. (**D**) XL-MS mapping of AHL protein homo- and heteromultimers. Crosslinked lysine residues were identified that sit immediately adjacent to the amino terminus of the Plants and Prokaryotes Conserved (PPC) domain and at the carboxyl end of the core sequence (GRP) of the type 1 AT-hook motif. The lysines are conserved across both homo- and heteromultimeric interactions, supporting a shared interaction interface.

We incorporated STRING confidence scores (28) into the network and color-coded the edges accordingly (Fig. 3A). These scores combine evidence from eight sources, including experimental data (Table S1) (28). For heteromeric PPIs, 28.2% (676) reached a threshold of ≥ 0.9, while 10% (241) fell within the 0.7–0.9 range. The remaining 1,468 interactions either scored below 0.7 or lacked prior annotation; this proportion increased further when considering only the experimental channel (Fig. 3C). These results suggest that our dataset captures both established and less-characterized PPIs.

For example, our data captured both homo- and heterodimeric associations among eight members of the AHL protein family (Fig. 3D). Previous yeast two-hybrid studies indicated that the PPC domain mediates these interactions (29). However, this hydrophobic domain often lacks lysine residues suitable for cross-linking (Fig. S8A-B). We identified conserved cross-linked lysine residues located adjacent to the N-terminus of the PPC domain and near the conserved GRP sequence within the AT-hook motif (Fig. 3D) (30). These cross-links provide direct evidence of spatial proximity within the complex. The cross-linked sites are conserved across both homo- and heterodimeric interactions, consistent with a shared interaction interface.

We also mapped the topologies of numerous well-characterized protein complexes (Figs. S9–S10), including the 26S proteasome; several chaperone and chaperone-like complexes, such as HSP70, the nascent polypeptide-associated complex, the TCP-1/cpn chaperonin (CCT), the prefoldin complex, and the 14-3-3 complex (GRFs); as well as several ATPases from chloroplasts, mitochondria, and vacuoles, in addition to various enzymatic complexes. Notably, AT2G07698, which encodes the ATPase F1 complex alpha subunit, and ATP-B (AT5G08670), both nuclear-encoded proteins, were found to form cross-links with chloroplast-encoded ATPase complex subunits as well as mitochondria-encoded ATPase subunits, suggesting dual localization and assembly in both organelles.

Together, STRING and PANTHER analyses indicate that systematic XL-MS captures both established and less characterized PPIs, complementing curated resources by identifying additional associations and providing structural context for their interaction interface.

### XL-MS provides structural insight into the topology of chloroplast 70S ribosome stalk regions inaccessible by cryo-EM

Extensive cross-links were identified for chloroplast 70S ribosomes, which are located on the stromal side of the chloroplast. In total, we identified 714 cross-links across 37 ribosomal subunits, including 616 intraprotein, 93 heteromeric, and 5 homomultimeric cross-links. Notably, many cross-links mapped to the flexible stalk regions of the chloroplast 70S ribosome (Fig. 4A–C, Fig. S11A–D), which have remained difficult to resolve because of the intrinsic dynamics of proteins such as uL1c, uL10c, uL11c, and bL12c (31, 32). A substantial proportion of the cross-links involved the flexible stalk protein bL12c, which forms the factor-recruiting arms of the ribosomal stalk (33). Despite its functional importance, bL12c remains poorly resolved in high-resolution structural studies because of its pronounced flexibility (34, 35); accordingly, it is absent from the high-resolution spinach 70S ribosome structure (PDB: 5MMM). By integrating XL-MS data with AlphaFold predictions (AF-P36210-F1-v6) and available PDB coordinates, we mapped the connectivity of bL12c within the ribosome. The identified cross-links revealed interactions between bL12c and other stalk subunits (uL10c and uL11c; green), the 50S subunit protein uL6c (red), and the 30S subunit proteins bS20c and uS12c (blue) (Fig. 4A).

**Figure 4.**
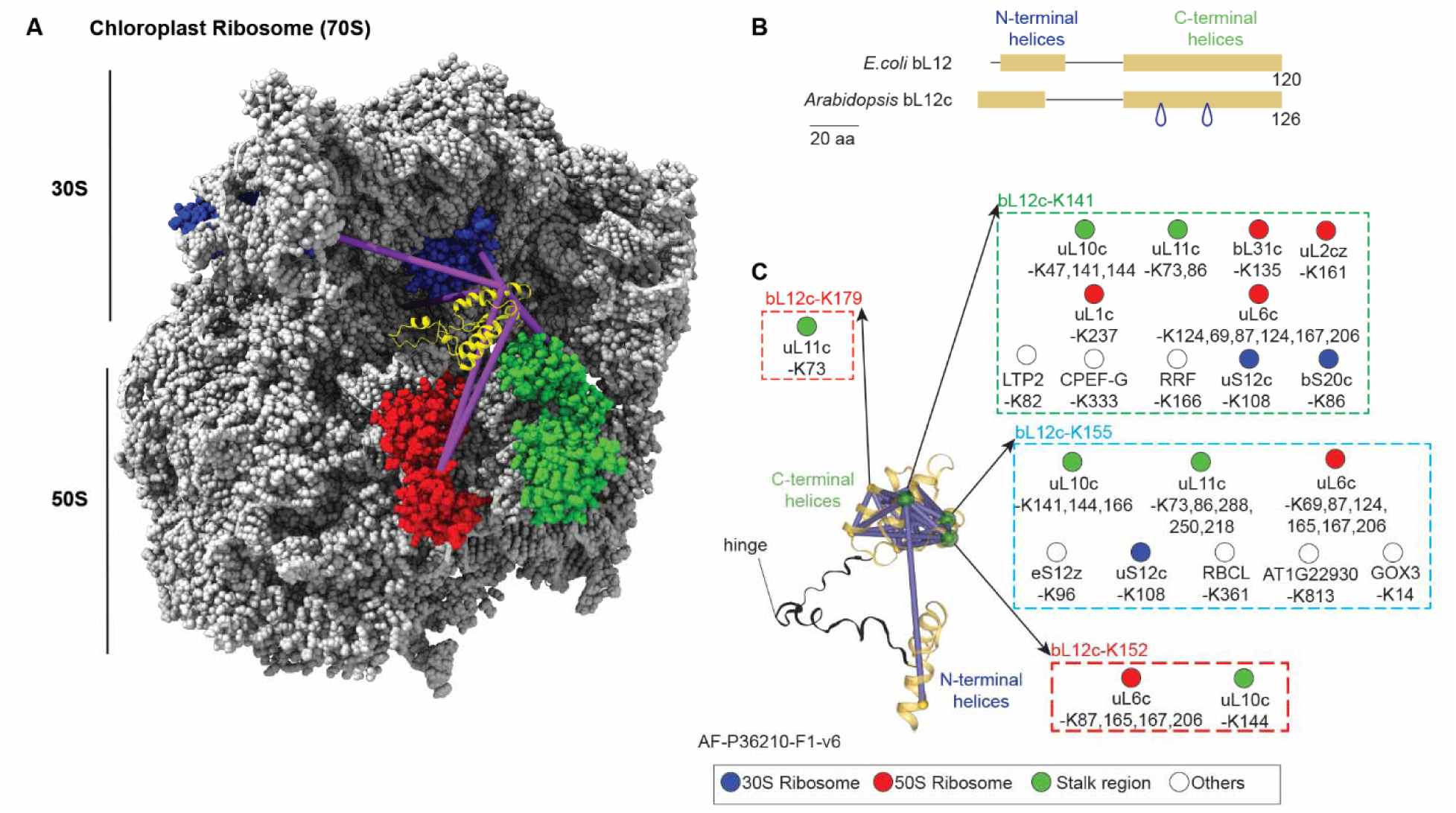
XL-MS provides structural insight into the topology of chloroplast 70S ribosome stalk regions inaccessible by cryo-EM. (**A**) bL12, a stalk region subunit, is not resolved in the cryo-EM structure of the spinach 70S ribosome (PDB: 5MMM). Integration of XL-MS data with AlphaFold predictions (AF-P36210-F1-v6) and PDB coordinates reveals its connectivity to other ribosomal components. Cross-links map interaction interfaces of bL12c with other stalk subunits (uL10c, uL11c, green), the 50S subunit (uL6c, red), and the 30S subunit (bS20c, uS12c, blue). The transit peptide sequence of bL12c was removed in the AlphaFold structure. (**B**) Domain comparison between *Arabidopsis* bL12c and *E. coli* bL12. Both homologs exhibit conserved N- and C-terminal helices connected by a flexible hinge region. XL-MS data identify homodimer interfaces localized to the C-terminus of bL12c. (**C**) XL-MS mapping onto the AlphaFold-predicted bL12c structure shows that lysine residues involved in inter-protein cross-links (K141, K152, K155, and K179) localize to the structured C-terminal domain. These cross-links suggest connections of bL12c not only with the 30S and 50S ribosomal subunits but also with the chloroplast elongation factor CPEF-G, the ribosome recycling factor (RRF), and other proteins. Nodes are color-coded by functional annotation category, and cross-linked sites are indicated.

While bL12c dimerization has been attributed to its N-terminal domain (NTD) (36), our data supported unexpected homodimeric cross-links within the C-terminal domain (CTD)—the region responsible for translation factor binding (Fig. 4B–C)—suggesting that CTD dimerization may provide a regulatory interface for partner recruitment. Additionally, these cross-links identify interactions between bL12c and chloroplast elongation factor CPEF-G, the ribosome recycling factor (RRF), and other proteins, underscoring the utility of XL-MS for dissecting the dynamic interactome of the ribosome.

### XL-MS identifies dynamic interaction networks in photosystem complexes

Photosystems I and II (PSI and PSII) and their associated light-harvesting complexes (LHCI and LHCII) are essential for light harvesting and carbon fixation but require continuous repair under fluctuating light conditions (37–39). LHCII forms PSII–LHCII supercomplexes and can transiently associate with PSI during state transitions, highlighting its dynamic nature and broad interaction potential. Although substantial progress has been made in understanding PSII repair (40, 41), many mechanistic details and repair intermediates remain unresolved.

Using XL-MS, we identified extensive interactions between PSII proteins and the repair factors PSB27, PSB28, PSB32, PSB33, and PSBTN (Fig. 5A). Unlike its cyanobacterial counterpart, which functions in both assembly and repair, Arabidopsis PSB27 appears to be specialized for repair, whereas the divergent PSB27-H1 (AT1G05385) has been implicated in assembly. Arabidopsis PSB27 lacks a lipid anchor and is structurally more flexible than the cyanobacterial protein (42) (Fig. S9A). A *Chlamydomonas* PSII structure (PDB: 6KAC), capturing PSB27 associated with both the PSII core and the oxygen-evolving complex (OEC), resolved residues 91–114 of PSB27 (43). Our XL-MS data showed that PSB27 interacts with both the PSII reaction center and OEC, involving K93 as well as K154 and K168 (Fig. 5A; Figs. S9B, S10A–H). These distance constraints support a more dynamic and flexible mode of PSII repair in plants (Fig. S9B). Comparison with pea cross-linking data (44) further identified several conserved PSB27 interaction sites (Fig. S13).

**Figure 5.**
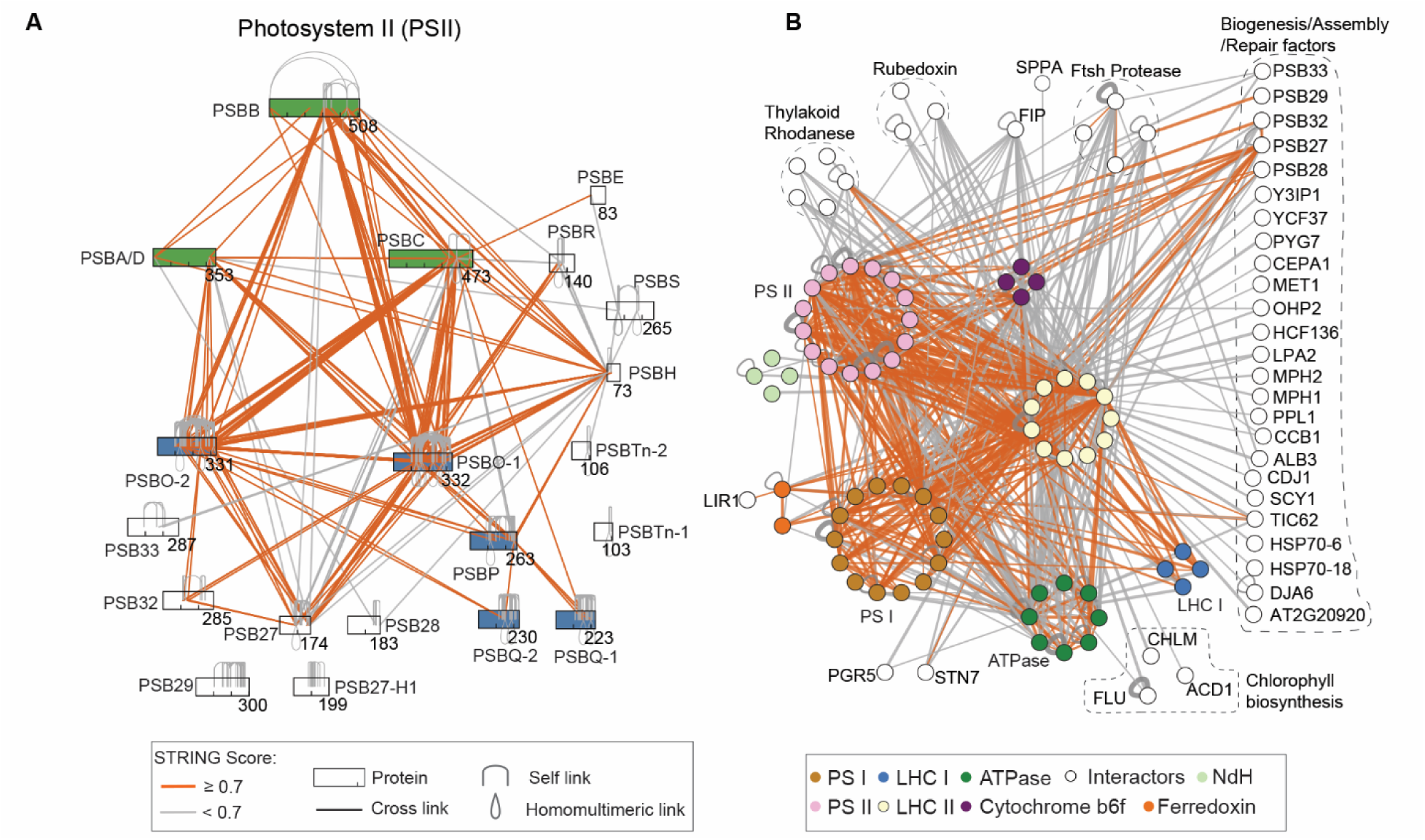
XL-MS identifies dynamic interaction networks in photosystem complexes. Cross-link edges are color-coded by STRING scores. (**A**) XL-MS identifies connectivity within a heterogeneous PSII population, mapping interactions among PSII subunits as well as with several repair and assembly factors, including PSB27, PSB28, PSB32, PSB33, and PSBTn-2. (**B**) XL-MS uncovers extensive interactions among PSI, PSII, LHCI, LHCII, ATPase, FtsH/SPPA proteases, and transient proteins involved in state transitions, assembly, repair, and chlorophyll metabolism.

XL-MS further revealed extensive connectivity among PSI, PSII, LHCI, LHCII, ATPase, FtsH/SPPA proteases, and transient factors involved in state transitions, assembly, repair, and chlorophyll metabolism. We detected interactions between STN7 kinase and LHCB1, LHCB2, and LHCB5, suggesting the dynamic remodeling of PSII–LHCII supercomplexes during state transitions and antenna disassembly during PSII repair (45, 46). In addition to its role in energy transfer, LHCII is associated with chlorophyll biosynthesis and catabolism factors, including ACD1, CHLM, and FLU, suggesting coordination of pigment homeostasis and ROS control during photosystem biogenesis. LHCII also interacted with multiple assembly and repair factors, including thylakoid rhodanese, MET1, OHP2, HCF136, LPA2, MPH2, and PPL1, consistent with a poised state that may facilitate rapid photosystem remodeling. Together, these findings support a model in which LHCII functions not only as a mobile antenna complex but also as an organizational hub linking photosystem assembly, repair, and environmental adaptation (45, 46).

### XL-MS of ribosome complexes reveals conserved 80S ribosome-associated factors and less characterized interactors

We identified extensive interactions involving the 80S ribosome, the complex composed of 60S and 40S subunits responsible for synthesizing nuclear-encoded proteins. Ribosomes assemble in the nucleolus, localize to the cytoplasm, and associate with the endoplasmic reticulum (ER). As a dynamic macromolecular machine, the ribosome coordinates translation through interactions with numerous proteins and RNAs. Perturbations in translation that lead to misfolded proteins, or inherent defects in the ribosome itself, can trigger specialized ribosome-associated quality control pathways (47). Using XL-MS, we mapped detailed structural information within this highly heterogeneous complex, capturing not only intra-and inter-subunit contacts (Fig. S14A-D), but also interactions with proteins involved in ribosome biogenesis, translation initiation, elongation and termination, as well as protein folding, quality control, stress response, and other regulatory pathways (Fig. 6A).

**Figure 6.**
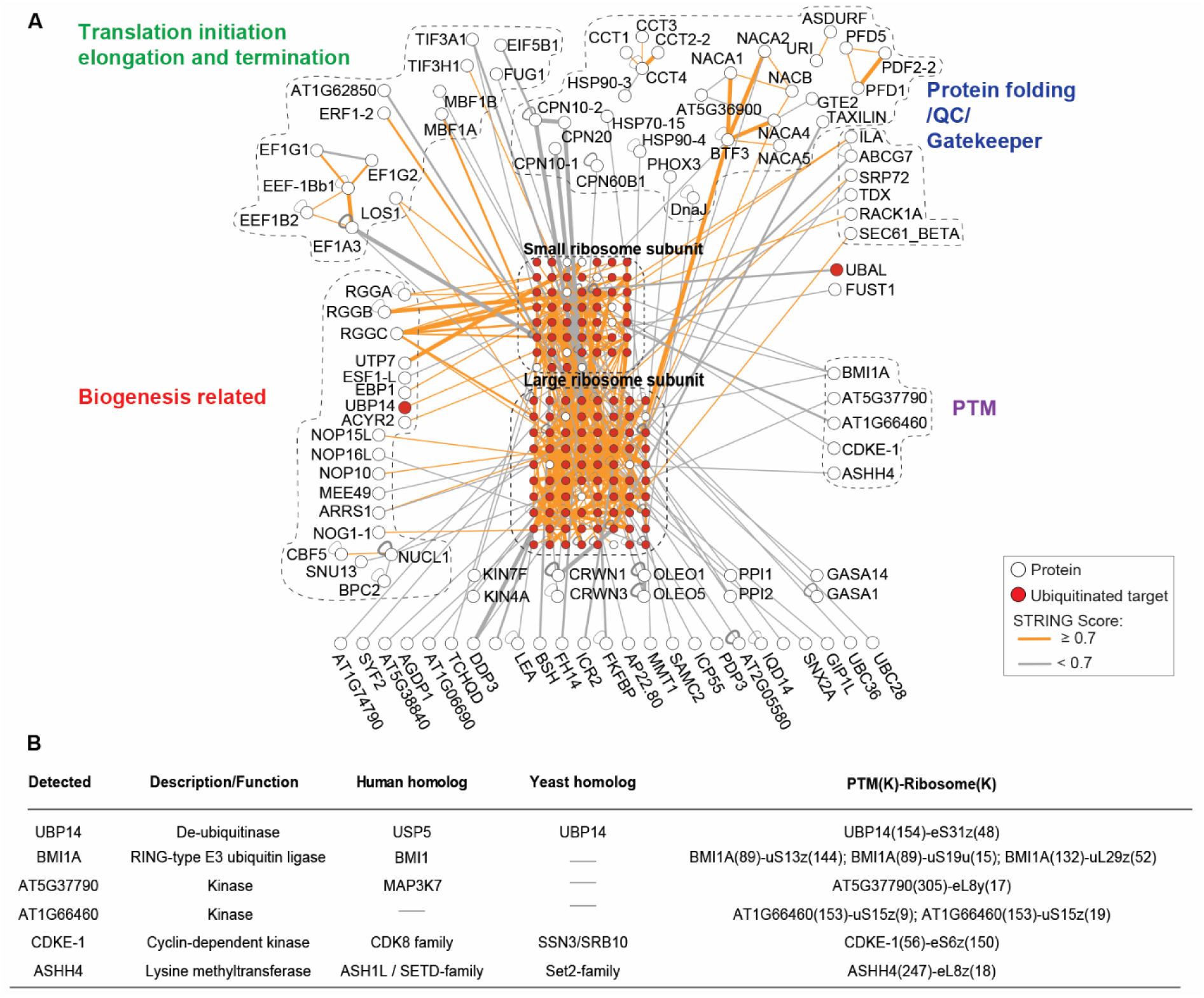
XL-MS of ribosome complexes reveals conserved 80S ribosome-associated factors and less characterized interactors. (**A**) XL-MS identified extensive interactions within 80S ribosomal subunits as well as with proteins involved in ribosome biogenesis, translation, folding, quality control, stress response, and other regulatory pathways. Ribosomal proteins are overlaid with ubiquitination data; red nodes indicate ubiquitinated proteins. (**B**) Table of XL-MS-identified ribosome-associated interactors with potential post-translational regulatory functions, many of which remain uncharacterized in plants. The description, human and yeast homolog by blast, and exact cross-linked sites are listed.

The functions of these interacting proteins in plants remain largely unknown despite extensive characterization in other organisms. Our XL-MS data suggest conserved roles for these candidates: UTP7 (AT3G10530) and NOG1-1 are likely involved in ribosome biogenesis (48, 49), whereas TAXILIN (AT5G50840) and ILITHYIA (ILA) may function in protein folding or translation (50, 51). RACK1A may modulate translation capacity (52, 53), while MBF1A and MBF1B may act in stress responses at stalled ribosomes, consistent with their roles in yeast (54). Additional candidates include RGG- and NOP-family ribosome biogenesis factors, as well as chaperones (e.g., HSPs, CPNs, tetratricopeptide repeat proteins such as PHOX3, and DNAJ proteins) implicated in protein folding and quality control (Fig. 6A). Notably, our analysis identified numerous less characterized interactions (gray edges), providing a resource for future studies of ribosome-associated factors in plants.

Ribosomes undergo extensive post-translational modifications (PTMs), including phosphorylation, methylation, and ubiquitination (55–59). Our XL-MS data identified several PTM-related factors associated with ribosomal proteins (Fig. 6B), including the deubiquitinase UBP14, the RING-type E3 ligase BMI1A, the lysine methyltransferase ASHH4, CDKE-1, and two additional kinases (AT5G37790 and AT1G66460). Notably, BMI1A formed cross-links with multiple ribosomal subunits, prompting us to further investigate ribosome-associated ubiquitination using diGly proteomics. Combined with targeted data mining, this analysis identified ubiquitination sites on 37 large-subunit and 27 small-subunit ribosomal proteins (Fig. 6A; Table S4), indicating widespread in vivo ubiquitination of ribosomal proteins in Arabidopsis.

### Histone interactome identifies established and less-characterized chromatin-associated interactions

Our XL-MS analysis identified extensive interactions between histones and a broad range of chromatin-associated proteins, providing insights into plant chromatin organization and gene regulation (Fig. 7A). Detected interactions included nucleosome assembly proteins (NAPs), chromosome structure maintenance proteins (SMCs), nuclear RNA polymerase components (NRPs), chromatin regulators, transcription-associated factors, and PAF complex subunits.

**Figure 7.**
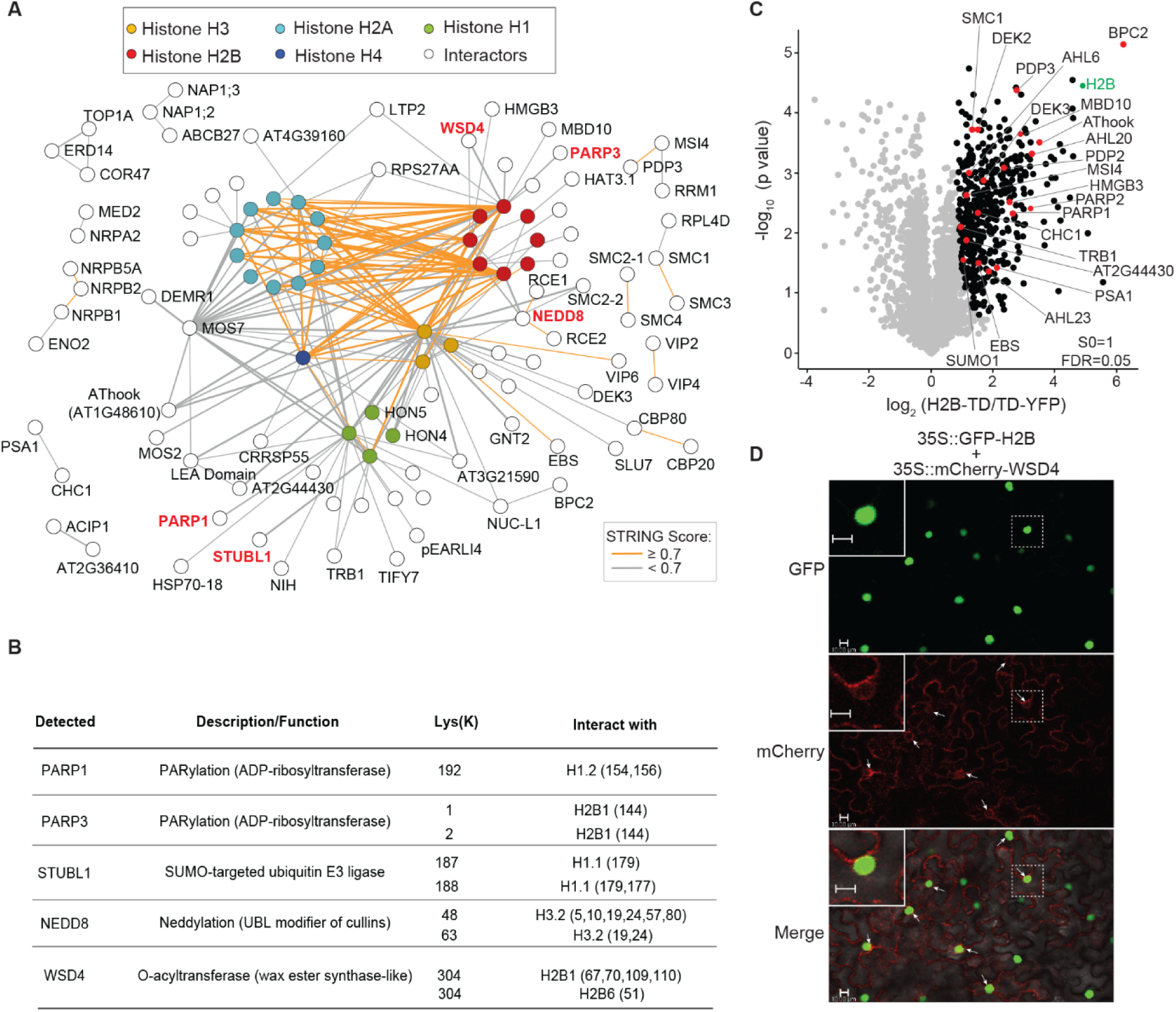
Histone interactome identifies established and less-characterized chromatin-associated interactions. (**A**) XL-MS reveals extensive interactions between histones and both known and previously uncharacterized proteins, including components of chromatin-associated complexes. Enzymatic regulators are highlighted in red. (**B**) XL-MS identified interactions between histones and PARP1, NEDD8, and WSD4, suggesting potential roles in histone modification and chromatin organization. Multiple cross-links between histone H2B proteins and WSD4, an O-acyltransferase, support a possible association of WSD4 with histone acylation-related processes. (**C**) Volcano plot of TurboID using H2B (AT3G45980) as bait, showing enrichment of proteins identified in proximity to histones. Nucleus-localized YFP-TD was used as the control. The x-axis represents fold enrichment, and the y-axis represents statistical significance (p-value). (**D**) Co-localization of GFP-HTB11 and mCherry-WSD4 in *Nicotiana benthamiana*. Green, GFP-HTB11; red, mCherry-WSD4. Overlapping nuclear signals are indicated by arrows, with enlarged views shown in the inset. Scale bar, 10 µm.

We further identified histone contacts with chromatin remodeling factors and epigenetic readers, including NUC-L1, EBS, DEK3, TRB1, bromodomain-containing proteins, MBD proteins, HMGB3, and the DNA helicase NIH. In addition, MOS7, a nucleoporin homologous to human NUP88, cross-linked with several histones, suggesting potential links between the nuclear envelope and chromatin organization (60). Histone interactions were also detected with regulatory proteins, including PARP1, PARP3, STUBL1, and the ubiquitin-like protein NEDD8. TurboID using H2B (AT3G45980) as bait supported the nuclear proximity of many identified candidates (Fig. 7B).

Surprisingly, XL-MS identified multiple distinct cross-links between histone H2B and the O-acyltransferase WSD4 (Fig. 7A, S12A-E). While WSD family members typically function in lipid metabolism and wax ester biosynthesis (61, 62), WSD4 is preferentially expressed in roots—tissues where wax deposition is uncommon, suggesting an alternative biological role (62). TAIR predicts a nuclear localization for WSD4, which we confirmed via co-expression of mCherry-WSD4 and GFP-H2B7 in *Nicotiana benthamiana*. They showed overlapping nuclear signals, as indicated by the arrows in Fig. 7B. (Fig. 7B). Together, these direct physical contacts and co-localization data suggest that WSD4 may catalyze histone acylation, a possibility that warrants further investigation (Fig. 7C).

Taken together, these findings expand the catalog of histone-associated proteins and enzymatic activities in plants, providing additional insights into chromatin biology and a resource for future studies of plant gene regulation.

## Discussion

Systematic mapping of PPIs is essential for understanding cellular function, yet plant interactomes remain less unexplored than those of other model organisms(63–66). This gap is significant given the importance of plant interactome studies for agriculture and environmental sustainability. Here, we established an XL-MS framework for mapping molecular machines and their interaction interfaces. To address the complexity of plant samples, we developed an optimized workflow integrating the PhoX cross-linker, which enables efficient IMAC enrichment of cross-linked peptides, together with subcellular fractionation and high-pH reversed-phase HPLC to increase proteome coverage.

Central to the workflow was pLink 3.2, which enabled efficient analysis of large-scale cross-linking datasets. Like other non-cleavable cross-linkers, however, PhoX-generated datasets remain computationally demanding because they involve a quadratic (n^2^) search space rather than simplified mass derivation through diagnostic ions used in MS-cleavable cross-linker (1). pLink 3.2 is specifically optimized to address this challenge and efficiently process large-scale datasets. In addition, because large-scale XL-MS is susceptible to FDR propagation, where low error rates at the CSM level can translate into substantially higher error rates at the PPI level (16, 17), we further validated FDR using an entrapment library search. Using this workflow with 1% FDR at the peptide-pair level, we identified 82,421 CSMs, corresponding to 52,944 cross-linked peptide pairs and 37,531 residue pairs across 5,064 proteins. These data supported the identification of 2,385 heteromeric PPI interactions, of which 51% were supported by multiple CSMs. Among the remaining 1,167 interactions supported by a single CSM, 215 had STRING confidence scores of at least 0.9, further supporting the reliability of the identified PPIs.

Current large-scale XL-MS studies have been performed in diverse systems, including cell lysates, isolated organelles, and intact cells (2). Although in vivo cross-linking best preserves the native cellular environment, its application in plants remains challenging because the cell wall limits cross-linker penetration and reduces peptide recovery. We therefore adopted a semi-native approach to achieve the depth and sensitivity required for systems-level analysis. We acknowledge that fractionation and lysis may introduce certain biases. For example, we detected an extensive interaction network associated with a specialized defense system in which the cytoplasm-localized GLL23 was crosslinked to the ER-body proteins BGLUs and JAL/PBP proteins (Fig. S17), likely reflecting interactions that occur upon cellular disruption (Fig. S17) (67, 68). These interactions are consistent with activation of the “mustard oil bomb” defense system and are therefore unlikely to represent random associations. We also observed good agreement with previous studies despite differences in isolation protocols; for instance, cross-links within the PSBO/PSB27/PSBC complex in Arabidopsis closely matched those reported from isolated pea chloroplasts (Fig. S13). In addition, a substantial proportion of the identified PPIs are supported by the STRING database or prior literature, indicating that our dataset captures biologically relevant protein complexes. Together, these data provide a representative snapshot of plant protein architectures and their interaction interfaces.

Many of these interactions were previously uncharacterized, particularly those involving ribosome-and histone-associated factors. For example, we identified WSD4, an O-acyltransferase, as a potential histone interactor. Multiple independent cross-links were detected between WSD4 and H2B (Fig. S15 and Fig. S16), and their association is further supported by nuclear co-localization. Although O-acyltransferases are primarily associated with lipid metabolism(61), their potential roles in histone modification remain largely unexplored. Given that several metabolic enzymes have been implicated in lysine acylation (69–72), our findings raise the possibility that WSD4 may contribute to chromatin regulation.

Previous studies have shown that heteromeric cross-links tend to exhibit larger median distances than homomultimeric cross-links, potentially reflecting greater conformational flexibility at heteromeric interfaces (73, 74). Accordingly, medium- and long-range cross-linkers may be better suited for protein–protein interaction mapping, whereas shorter cross-linkers may provide higher precision for structural modeling. PhoX contains a spacer arm of ∼5 Å (2, 6). Its overall arm length is longer than EDC (0 Å), but shorter than those of the MS-cleavable cross-linkers DSSO (10.1 Å), DSBSO (12.9 Å), and CBDPS (14.0 Å), and substantially shorter than that of PIR (29.3 Å). Reported distance constraints for PhoX range from 20 Å (11) to 35 Å (12); here, we adopted a 35 Å threshold to account for protein backbone flexibility and side-chain dynamics. Successful mapping of both intra- and inter-protein cross-links onto existing structures, together with extensive PPI identification, shows that PhoX is suitable for both structural analysis and interactome mapping. Future efforts toward comprehensive protein complex mapping will likely benefit from combining MS-cleavable and non-cleavable cross-linkers, using linkers with varying spacer lengths, and integrating complementary search engines. For example, comparison with xiSEARCH revealed a substantial overlap in identified cross-linked peptides, while each search engine also detected unique peptide pairs (Fig. S16A).

In summary, our results demonstrate that XL-MS can effectively map PPIs under semi-native conditions. By capturing both established and previously uncharacterized interactions, this study advances our understanding of the plant interactome and provides a valuable resource for investigating protein complex architecture. In addition, the extensive intra-protein cross-links offer structural constraints for protein modeling. Together, these data establish a framework and resource for exploring plant molecular machines that can be extended to other plant species and crops.

## Acknowledgement

This work was supported by the National Institutes of Health (NIGMS R01GM135706 and S10OD030441 to S-L.X.) and by the Carnegie endowment fund for the Carnegie Mass Spectrometry Facility. We thank Prof. Laura Helen Gunn for providing the structure of Arabidopsis Rubisco L8S8, and Drs. Mike Trnka and Robert Chalkley (UCSF) and Dr. Rosa Viner (Thermo Fisher Scientific) for technical assistance. We also thank the Hong Qiao lab (UT Austin) for the nuclear isolation protocol and the Adrien Burlacot lab for the chloroplast isolation protocol. We thank Dr. Ajeet Chaudhary and Prof. Arthur Grossman for their critical reading of the manuscript.

## Author contributions

S.L.X conceived the project and supervised the study. C.S.T. and R.S. designed and performed most of the XL-MS experiments. R.S. developed the initial XL workflow. W.C.C performed chloroplast XL-MS. S.K. performed ubiquitination enrichment. C.S.T and S.L.X analyzed all the MS data and prepared figures, with assistance from A.V.R, A.Y., G.L. and K.H. C.S.T and S.L.X interpreted the data and wrote the manuscript with feedback from all authors.

## Data Availability

The mass spectrometry proteomics data have been deposited to the ProteomeXchange Consortium via the PRIDE partner repository with the dataset accession numbers PXD066234, PXD066291, PXD073734 and PXD078284. All other data is available from the corresponding author on reasonable request.

## Supplemental Table

Table S1: Summary of cross-linked peptide, including intra- and inter- protein cross-links, PPI annotations based on STRING and FDR reports.

Table S2: SUBA5 annotation and Analysis of protein classification using PANTHER.

Table S3: mapping of intra-cross-links to known structures from PDB and predicted structure from AlphaFold, and simulation analysis comparison.

Table S4: Detected cross-links for ribosomal and histones PPIs, and ribosome-related ubiquitination.

Table S5: Names, accession numbers and description used in the figures and a glossary.

## Material and methods

### Plant growth condition

Arabidopsis thaliana wild-type (Col-0) seeds were sterilized and sown on ½ Murashige and Skoog (MS) agar square plates supplemented with 0.6% (w/v) phytoblend and lined with nylon fabric. After stratification at 4°C for 3 days, the plates were transferred to growth conditions.

Light-grown seedlings: plants were vertically grown in a growth chamber at 22°C under a 16-hour light / 8-hour dark cycle for 10-12 days, then flash frozen in liquid nitrogen and stored at -80°C for crosslinking experiments on cell lysates and isolated chloroplasts. Dark-grown seedlings: plants were grown horizontally at 22°C in complete darkness for 3 days in a growth chamber, then flash frozen in liquid nitrogen and stored at -80°C for nucleus isolation. Dark growth was used to reduce Rubisco content.

Frozen Arabidopsis tissue was cryo-ground to a fine powder. Different amounts of tissue were used for cross-linking experiments: 6 g from 10 d seedlings for cell lysates, 195 g from 12d seedlings for chloroplasts, and 32 g (replicate 1 with PhoX) or 14 g (replicate 2 with tbu-PhoX) from 3d dark-grown seedlings for nuclei.

### Fraction of plant seedling tissues for cell lysate, chloroplast and nucleus

#### Cell lysates

Homogenized 10 d old light growth tissues were extracted using MOPS buffer [100 mM MOPS pH 7.6, 150 mM NaCl, 1% Nonidet P-40, 1 mM PMSF, 1x Roche protease inhibitor] at a tissue:buffer ratio of 1:1 (g/mL, w/v). The samples were centrifuged at 20,000 g for 15 min at 4°C, and the supernatant was then filtered twice with Miracloth and used for the cross-linking reaction.

#### Chloroplast isolation

Frozen, homogenized tissue powder from 12 d old seedlings was mixed with a grinding buffer similar to (75), using HEPES instead of Tricine for crosslinker compatibility. Briefly, 5 g of tissue powder was ground in a mortar and pestle with 30ml of grinding buffer [20mM HEPES-KOH pH 8.0, 330mM sorbitol, 5 mM EDTA, 5 mM EGTA, 5 mM MgCl_2_, and 10 mM NaHCO_3_] at 4°C, and under green illumination only. The resuspended homogenate was filtered twice through two layers of Miracloth. 30 ml of the filtrate was carefully transferred onto 10 ml of 40% Percoll [20 mM HEPES-KOH pH 8.0, 330 mM sorbitol, 40% v/v Percoll, 2.5 mM EDTA, 5 mM MgCl_2_, and 10 mM NaHCO_3_] in a 50ml centrifuge tube, and centrifuged at 2,600g for 10 min at 4°C. The bottom pellet, enriched in intact chloroplasts, was collected and resuspended in 30 ml of wash buffer [20 mM HEPES-KOH pH 8.0, 330 mM sorbitol, 2.5 mM EDTA, 2 mM MgCl_2_, and 10 mM NaHCO_3_], and centrifuged at 2,600g for 3 min at 4°C. Each pellet derived from 5 g of tissue was washed twice with 4.5 ml of 1x PBS supplemented with MgCl_2_ [137 mM NaCl, 2.7 mM KCl, 10 mM Na_2_HPO_4_, 1.8 mM KH_2_PO_4_, and 2.5 mM MgCl_2_] and then resuspended in 1mL 1x PBS supplemented with MgCl_2_ for the cross-linking reaction. This process was repeated across multiple batches, with a total of 195 g of tissue used for cross-linking.

#### Nuclear isolation

The method for nuclear isolation was adapted from (76) with slight modification. Powdered dark-growth Arabidopsis tissue of approximately 2 g was completely homogenized with lysis buffer [20 mM HEPES (pH 7.4), 20 mM KCl, 2 mM EDTA, 2.5 mM MgCl_2_, 25% glycerol, 250 mM sucrose, 1 mM dithiothreitol, 1 mM phenylmethylsulfonyl fluoride (PMSF), 1× protease inhibitor cocktail, and 0.2% formaldehyde] at tissue:buffer ratio of 1:5 (g/mL, w/v). The homogenate was then filtered through a 40 μm cell strainer and centrifuged at 4°C, 1,500g for 10 min. The pellet was retained as the nuclear fraction. The nuclear pellet was then washed 5 times with 5 mL of nuclear washing buffer [20 mM HEPES (pH 7.4), 2.5 mM MgCl_2_, 25% glycerol, 0.2% Triton X-100, 1 mM PMSF, and 1× protease inhibitor cocktail] and centrifuged at 4°C, 1,500g for 10 min to collect the pellets after each wash. The washed nuclear pellet was then resuspended in 0.5 mL of nuclear resuspension buffer 1 [20 mM HEPES (pH 7.4), 10 mM MgCl_2_, 250 mM sucrose, 0.2% Triton X-100, 1 mM PMSF, and 1× protease inhibitor cocktail] and carefully laid on top of nuclear resuspension buffer 2 [20 mM HEPES (pH 7.4), 10 mM MgCl_2_, 1.7 M sucrose, 0.2% Triton X-100, 1 mM PMSF, and 1× protease inhibitor cocktail], then centrifuged at 4°C,16,000g for 45 min. The supernatant was carefully removed, and the nuclear pellet was resuspended in 0.5 mL 1 X PBS (pH 7.5) buffer for crosslinking reaction. The intactness and purity of the nuclear pellet was assessed by DAPI staining under a Leica SP8 confocal microscope. Multiple samples were prepared in parallel, and final isolated nuclei were pooled before crosslinking.

### Cross-linking reaction

PhoX or tBU-PhoX cross-linker was freshly prepared at a stock concentration of 100 mM in DMSO for each cross-linking experiment. A working concentration of 2-5 mM cross-linker was added to the buffer solution containing whole cell lysate, isolated chloroplasts or isolated nuclei and incubated for 2 h at 4°C (dark condition for chloroplast cross-linking) with end-to-end rotation. The cross-linking reaction was quenched by adding Tris pH 8.0 to a final concentration of 20 mM and subjected to protein extraction.

### Protein extraction, digestion, IMAC enrichment and RPLC fractionation

Protein extraction from the cross-linked samples, digestion to peptides, immobilized metal affinity chromatography (IMAC) enrichment, and high pH reverse HPLC fractionation followed the same procedure as described in (13). Multiple batches of PhoX-cross-linked peptides from isolated intact chloroplasts were combined before IMAC enrichment. More information can be found in supplementary Fig. S3. Multiple batches of samples were combined prior to high pH reversed-phase HPLC offline fractionation (RPLC) on Vanquish Flex HPLC, and fractions were collected for further LC-MS/MS analysis.

To generate selected proteomic libraries, total peptides from cell lysate, chloroplast and nucleus samples were fractionated into multiple fractions using the Pierce High pH Reverse-Phase Peptide Fractionation Kit and were analyzed by LC-MS/MS.

### diGly proteomics analysis to identify ubiquitinated proteins *in vivo* and datamining

Metabolically labeled Arabidopsis plants with ^14^N and ^15^N isotope were harvested, followed by protein extraction and trypsin digestion as previously described (13, 77). Ubiquitinated peptides were enriched using the PTMScan® Ubiquitin Remnant Motif (K-ε-GG) Kit (Cell Signaling Technology) according to the manufacturer’s instructions. The enriched peptides were then analyzed by LC-MS/MS. Peptides identified in both light and heavy forms were considered high confidence. Ubiquitination sites were further identified by data mining as described in (78).

### LC-MS/MS analysis

Cross-linked peptides were analyzed by liquid chromatography-tandem mass spectrometry (LC-MS/MS) using an Easy-nLC 1200 ultra-performance liquid chromatography (UPLC) system (Thermo Fisher) coupled to an Orbitrap Eclipse Tribrid quadrupole Orbitrap mass spectrometer (Thermo Fisher). Peptides were first loaded onto a C18 trap column (Acclaim PepMap 100 C18) and then separated on an analytical C18 column (Aurora Series, 25 cm × 75 µm ID, Ion Opticks). The flow rate was set to 300 nL/min and the separation was performed over a 120-minute gradient. Peptides were eluted using a gradient from 3% to 28% solvent B over 106 minutes, followed by an increase from 28% to 44% over 15 minutes and a final wash at 90% solvent B for 14 minutes. Solvent A consisted of 0.1% formic acid and solvent B consisted of 80% (v/v) acetonitrile with 0.1% (v/v) formic acid.

Data was acquired on an Orbitrap Eclipse mass spectrometer. Full MS scans were acquired over a mass-to-charge (m/z) range of 375–1600 at a resolution of 120,000, with an automatic gain control (AGC) target of 200,000, a maximum injection time of 50 ms, normalized AGC target set to 50%, and RF lens set to 30%. The most intense multiply charged precursors (charge states 2–8) were selected for fragmentation with a cycle time of 3 seconds. MS/MS scans were acquired at a resolution of 15,000 with an AGC target of 5 × 10⁴, maximum injection time of 22 ms, isolation window of 1.4 m/z, and normalized AGC target set to 100%. Fragmentation was performed using higher-energy collisional dissociation (HCD) with a normalized collision energy (NCE) of 27. Dynamic exclusion was enabled for 30 seconds. For cross-linked peptide detection, charge states 3 to 8 were used.

For Real-time library search (RTLS) acquisition (15), a low-resolution MS2 survey scan performed in the ion trap over an m/z range of 120–500 was included. The resulting MS2 peaks were matched against a custom spectral library containing signature peak patterns characteristic of mono-links and cross-links (signature m/z values: 201.1231, 215.1387, 312.0632, and 377.1269, each with characteristic relative intensities for mono- or cross-linked species). Only precursor ions whose survey MS2 spectra matched the cross-link pattern using the “Similarity Search” function were selected for high-resolution MS2 acquisition.

### Data search using pLINK3.2

Linear peptide were searched against Arabidopsis TAIR database using Protein Prospector as described in (77). Phosphopeptide identifications were performed using the FragPipe pipeline(79). For all searches, Precursor and fragment mass tolerance were set up as 10 and 20 ppm.

Cross-linked peptide identification was performed using pLink3.2 (unreleased), an extension of the pLink 3 framework. Compared to pLink2 (14), pLink3 incorporates machine learning-based scoring, an optimized indexing structure, and improved FDR control (https://github.com/pFindStudio/pLink3), resulting in enhanced speed, sensitivity and precision. Version 3.2 further introduces a hierarchical funnel architecture that unifies searches for non-cleavable and cleavable cross-linkers (Mao et al, unpublished).

Search parameters were as follows: precursor and fragment mass tolerances of 5 ppm and 20 ppm, respectively; PhoX as the cross-linker (cross-linked mass 209.972 Da; mono-linked mass 227.982 Da), targeting Lys residues and protein N-termini; Trypsin/P digestion (allowing cleavage before Proline) with up to three missed cleavages; peptide length range of 6–60 amino acids and mass range of 600–6000 Da. Carbamidomethylation (Cys) was set as a fixed modification, while oxidation (Met) and protein N-terminal acetylation were specified as variable modifications.

The data were searched against specific databases for cell lysate (11,734 entries), chloroplast (9,798 entries), and nucleus (14,643 entries) using pLink 3.2. Results were filtered to a 1% false discovery rate (FDR) at the peptide-pair level. FDRs for mono-links, loop-links, and intra- and inter-protein cross-links were calculated independently. To compare two acquisition methods (XL-Standard [Standard] and Real-time library search [RTLS]), data acquired from the whole-cell lysate XL samples were used for evaluation.

FDR estimation in pLink 3 followed a three-step framework. First, results were aggregated at the peptide-pair level, grouping spectra by unique peptide backbone peptide backbone sequence and cross-linking sites, regardless of charge states and variable modifications. Second, FDR was calculated using target–decoy competition (FDR = (TD − DD) / TT). Third, empirical post-filters were applied to remove low-confidence identifications, such as PPIs supported by a single low-scoring CSM. At the residue-pair level, cross-linked peptides sharing identical linkage sites (including those arising from missed cleavages) were grouped, and these residue pairs were subsequently aggregated to the PPI level.

Entrapment-estimated FDP (false discovery proportion): For entrapment analysis(18), the data were searched against the Arabidopsis database supplemented with an equal number of human protein entries. To minimize false positives from conserved sequences, homologous human proteins were first identified by linear peptide searches (including regular, mono-, and loop-linked peptides) against the full human database and subsequently excluded. From the remaining pool, a subset of human proteins matched for database size and tryptic peptide distribution (K/R content) was selected and concatenated with the target database. FDRs for mono-links, loop-links, and intra- and inter-protein cross-links were calculated separately using pLink 3.2. FDPs were 0.4% (chloroplast), 2.7% (cell lysate), and 2.8% (nucleus) at the peptide-pair level, and 1.8%, 4.4%, and 5.7% at the PPI level, respectively.

### Data compilation and reporting

Intra- and inter-protein cross-links were compiled from multiple experiments (Supplementary Fig. S3) and reported separately (Table S1). A CSM was defined as distinct based on differences in peptide backbone sequence, cross-linking site, charge state, or variable modifications (e.g., oxidation). A peptide-pair was defined as distinct on differences in peptide backbone sequence and cross-linking site regardless of charge state or modification. Residue pairs were defined by the cross-linked sites, grouping peptides with identical linkage positions, including those arising from missed cleavages.

### Data search using xiSEARCH

MGF files for the xiSEARCH were generated using MSConvert (ProteoWizard v3.0.26049) following Xi guidelines. Searches performed with xiSEARCH v1.8.13 and xiFDR v2.3.10 using parameters aligned with pLink 3.2. The boost was set to the peptide-pair level, and the boost "between" option was enabled. All other parameters in xiSEARCH and xiFDR were kept at their default settings.

### Subcellular localization analysis of proteins using SUBA5

The subcellular localization of cross-linked proteins was analyzed using SUBA5 (https://suba.live/). Accession IDs of proteins identified in both intra- and inter-protein cross-links were submitted as input. Only the localization consensus data were used for classification. Proteins assigned to multiple compartments were counted once in each corresponding localization category when calculating total counts.

### PANTHER protein class analysis and mapping intralinks to PDB and AlphaFold structures using ComMap

Arabidopsis gene identifiers (ATG accession numbers) were used for protein class analysis using PANTHER™ Protein Class tool (27), which contains 210 classification terms (version 19.0, released 2024-06-20).

To map the results of intralinks against experimentally determined structures from the Protein Data Bank (PDB) and predicted structures from AlphaFold protein structure database, the protein accession IDs were first converted to UniProt IDs. The intralink information was then mapped to available structures using ComMap, a tool that enables large-scale structure-based mapping by integrating XL-MS data with existing structural models and performing distance calculations (19).

### Crosslinking data visualization

#### Cytoscape visualization

The entire cross-link-based protein-protein interaction (PPI) network was visualized in Cytoscape (version 3.10, https://cytoscape.org/) using inter-protein cross-link data, including heteromeric and homomultimeric cross-links. Nodes were colored according to protein class information from the PANTHER classification system, and edges were colored according to the STRING interaction score (https://string-db.org/).

#### xiVIEW

PPIs, including both inter- and intra-protein cross-links, were visualized using xiVIEW (https://xiview.org/) (80). PPIs were categorized via the built-in STRING data loader (score cutoff 0.4).

Cross-linked peptides supporting the same lysine pair were consolidated into a unique K-K linkage, collapsing redundancy (e.g. due to miscleavage events) and reporting each linkage only once.

#### Mapping of crosslinks to high-resolution structures of proteins/complexes

Intra- and inter-crosslinks were superimposed on PDB or AlphaFold structures using the built-in function of xiVIEW, which also calculated the corresponding crosslink distances within the high-resolution structures. Alignments of protein orthologs were performed and visualized using PyMOL (https://www.pymol.org/).

### Histone-TurboID Proximity-labeling

Histone H2B (H2B6, AT3G45980) was cloned into the Gateway system and recombined into the destination vector pB7m34GW to generate the ACINUSpro::H2B6-TurboID-VENUS construct, which was introduced into *Arabidopsis* wild-type plants via floral dipping. T2 transgenic seedlings were used for TurboID experiments. Plants expressing ACINUSpro::TurboID-YFP-NLS served as controls. TurboID sample preparation, data acquisition, and analysis were performed as previously described (23, 81).

### Co-localization assay

Co-localization was assessed in *Nicotiana Benthamian* leaves using transient co-expression. WSD4 was cloned into a modified pMDC43 Gateway binary vector containing an N-terminal mCherry tag to generate 35S::mCherry-WSD4, and H2B7 was introduced to pGWB6 gateway vector to generate 35S::sGFP-H2B7. Agrobacterium-mediated infiltration was performed, and fluorescence signals were visualized three days post-infiltration using a Leica SP8 confocal microscope.

---

**Supplementary Fig. S1:**
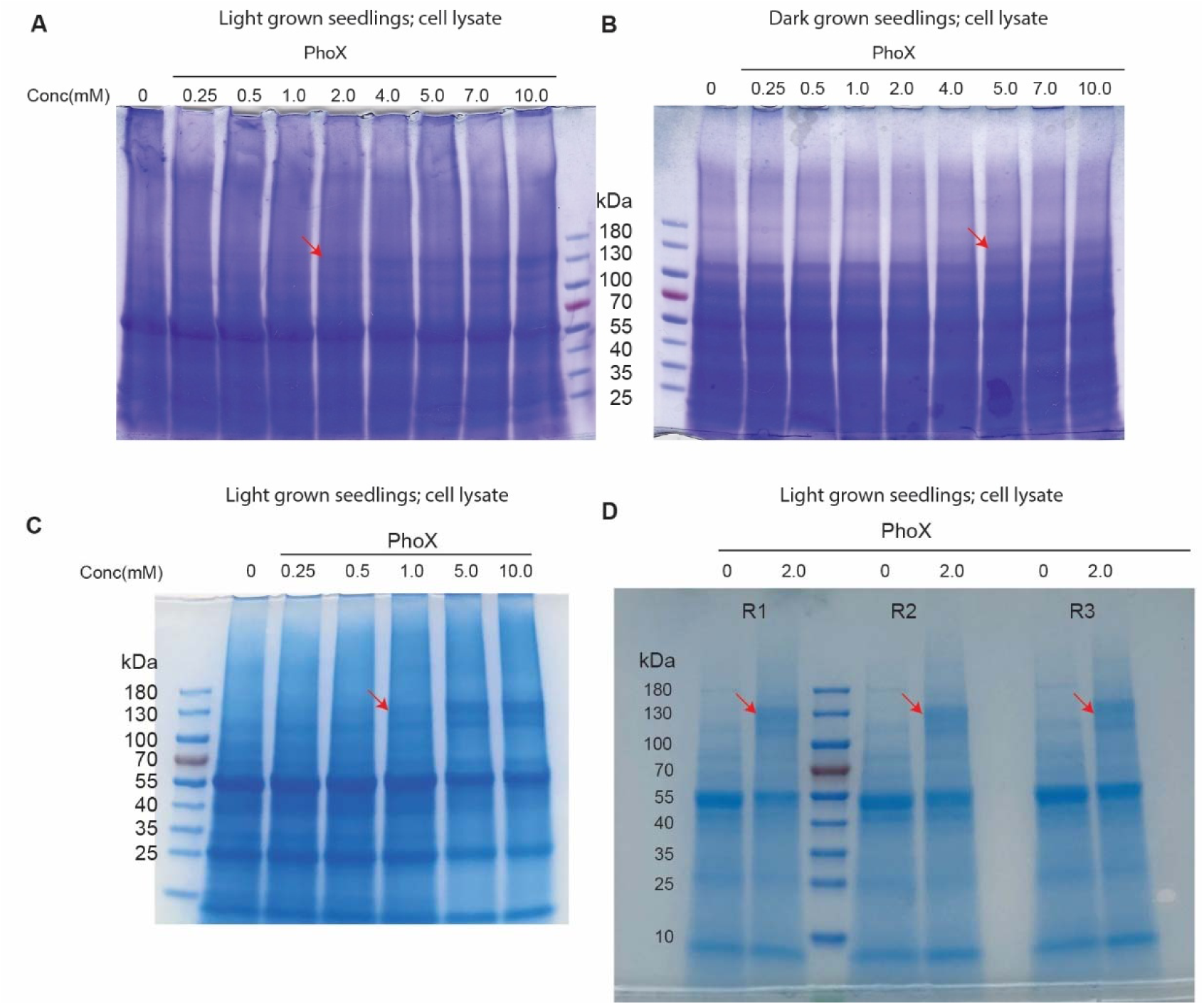
Optimization of the PhoX cross-linker concentration for XL-MS. (**A**-**D**) Different concentrations of PhoX cross-linker (0 to 10 mM) were tested on cell lysates from light (A, C, D) or dark grown seedlings (B). Increased protein cross-linking, indicated by the accumulation of higher molecular weight proteins (red arrow), was observed with increasing cross-linker concentration. However, no additional visible cross-linking was detected at higher concentrations (7-10 mM).

**Supplementary Fig. S2:**
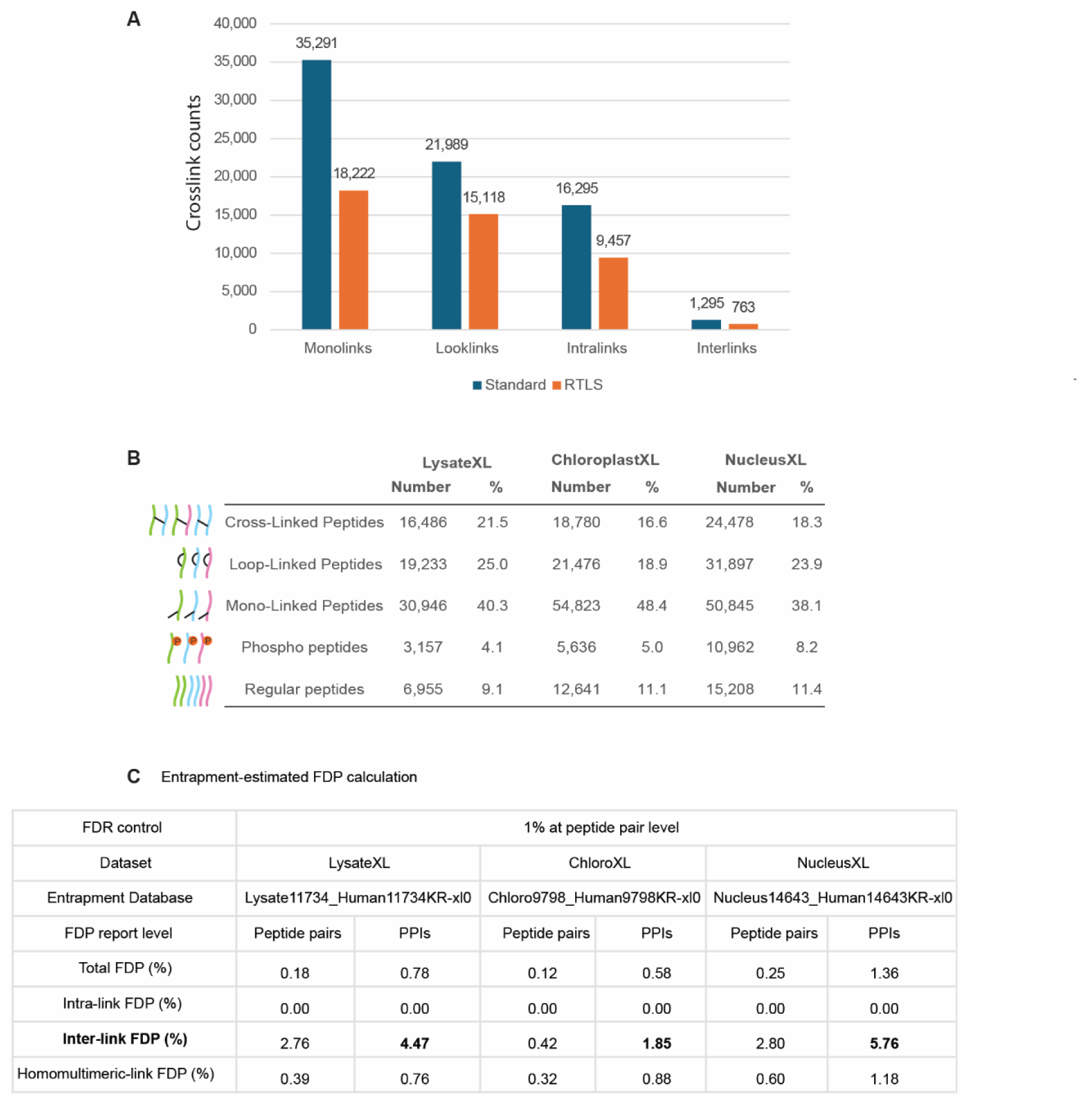
Acquisition method comparison and IMAC enrichment evaluation. **(A)** Comparison of acquisition strategies shows that standard XL acquisition outperforms Real-Time Library Search (RTLS). Data were acquired on an Orbitrap Eclipse mass spectrometer using PhoX-enriched cross-linked peptides from *Arabidopsis* cell lysates and searched against the lysate proteome protein database. RTLS acquisition reduced the identification of monolinks, defined as species in which one NHS ester reacts with a peptide while the other is hydrolyzed, by approximately 48%, and loop-links, defined as cross-links formed within a single peptide, by about 31%. However, RTLS also markedly decreased the detection of inter peptide cross-links, including both intra-protein (intralinks) and inter-protein (interlinks) interactions, by 42% and 41%, respectively. **(B)** Percentage analysis of detected peptides indicates efficient enrichment using the IMAC workflow, with regular non phospho-modified linear peptides comprising less than 12% of all identifications. Notably, because precursor ions with charge states of 3+ or higher were preferentially selected for MS/MS analysis, the true proportions of both regular peptides and phosphopeptides may be slightly underestimated. For simplicity, only the peptide backbone sequence, excluding PTM modifications and cross-linking sites, was considered, and unique peptide counts are reported. The relative abundance trends, from most to least abundant, are consistent across cell lysate, chloroplast, and nucleus samples. All data were derived from pLink 3.2 searches, except for phosphopeptide identifications, which were obtained using FragPipe. **(C)** False discovery proportion (FDP) analysis at the peptide-pair and PPI levels was performed using a human entrapment library to estimate the FDP. Separate entrapment databases were generated for each dataset and searched independently using pLink 3.2.

**Supplementary Fig. S3:**
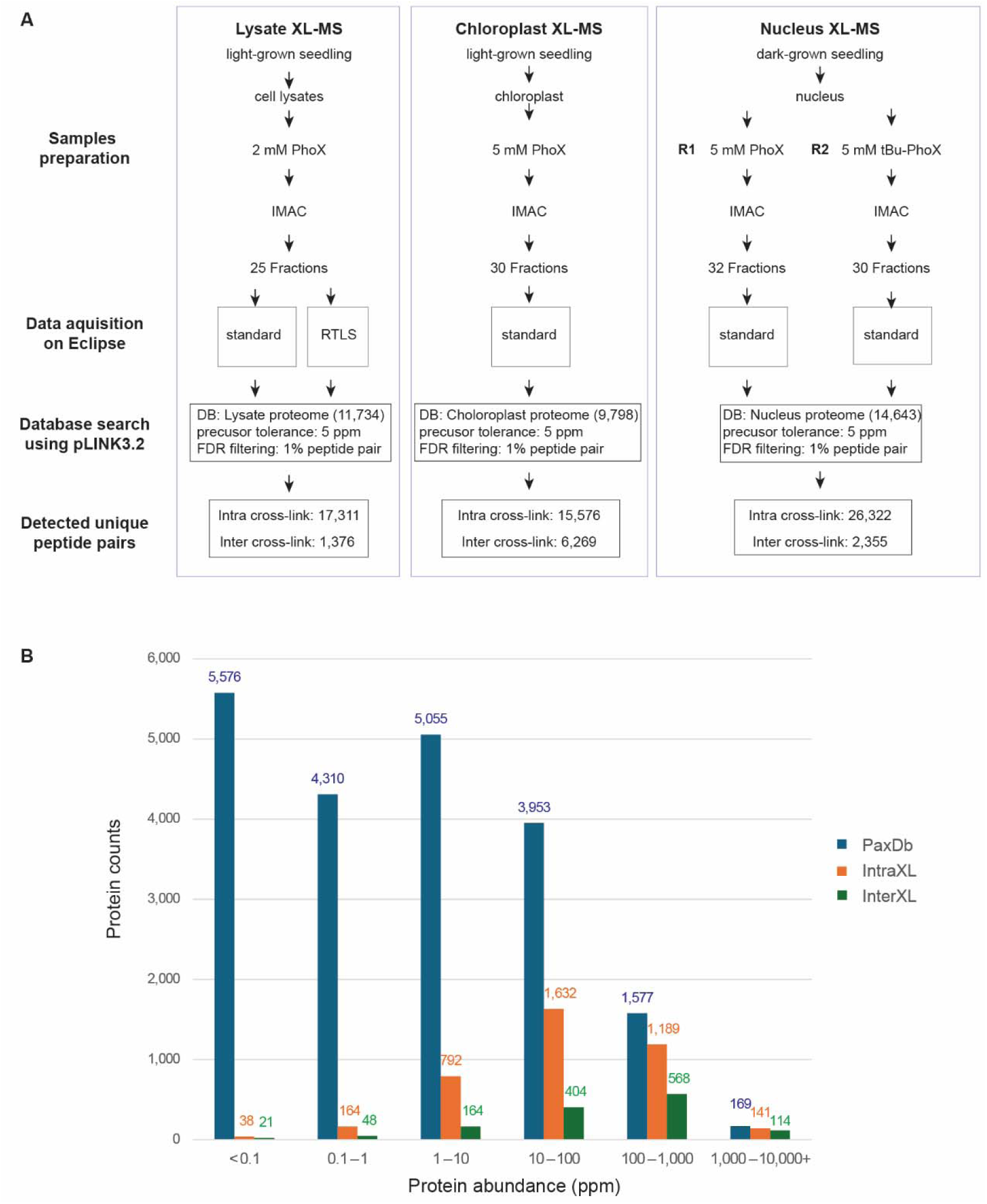
Summary of XL-MS experiments. (**A**) Key details of sample preparation, data acquisition using the Orbitrap Eclipse mass spectrometer, and database searching using pLINK3.2 are provided. A summary of the unique cross-linked peptide pairs identified across experiments is also included. The peptide-pair is defined by distinct peptide backbone sequence, and cross-linking sites, regardless of charge states and variable modifications. (**B**) The distribution of cross-linked protein identifications across abundance ranks, based on the PaxDb index (pdx-dp.org; blue), indicates broad proteome coverage. In addition to highly abundant proteins, a substantial number of medium- and low-abundance proteins were also identified as cross-linked for both intra-protein (orange) and inter-protein (green) crosslinks, highlighting the depth of the approach. The x-axis represents protein abundance, expressed in ppm (parts per million), reflecting the relative abundance of each protein compared to all others in the sample, while the y-axis shows the counts.

**Supplementary Fig. S4.**
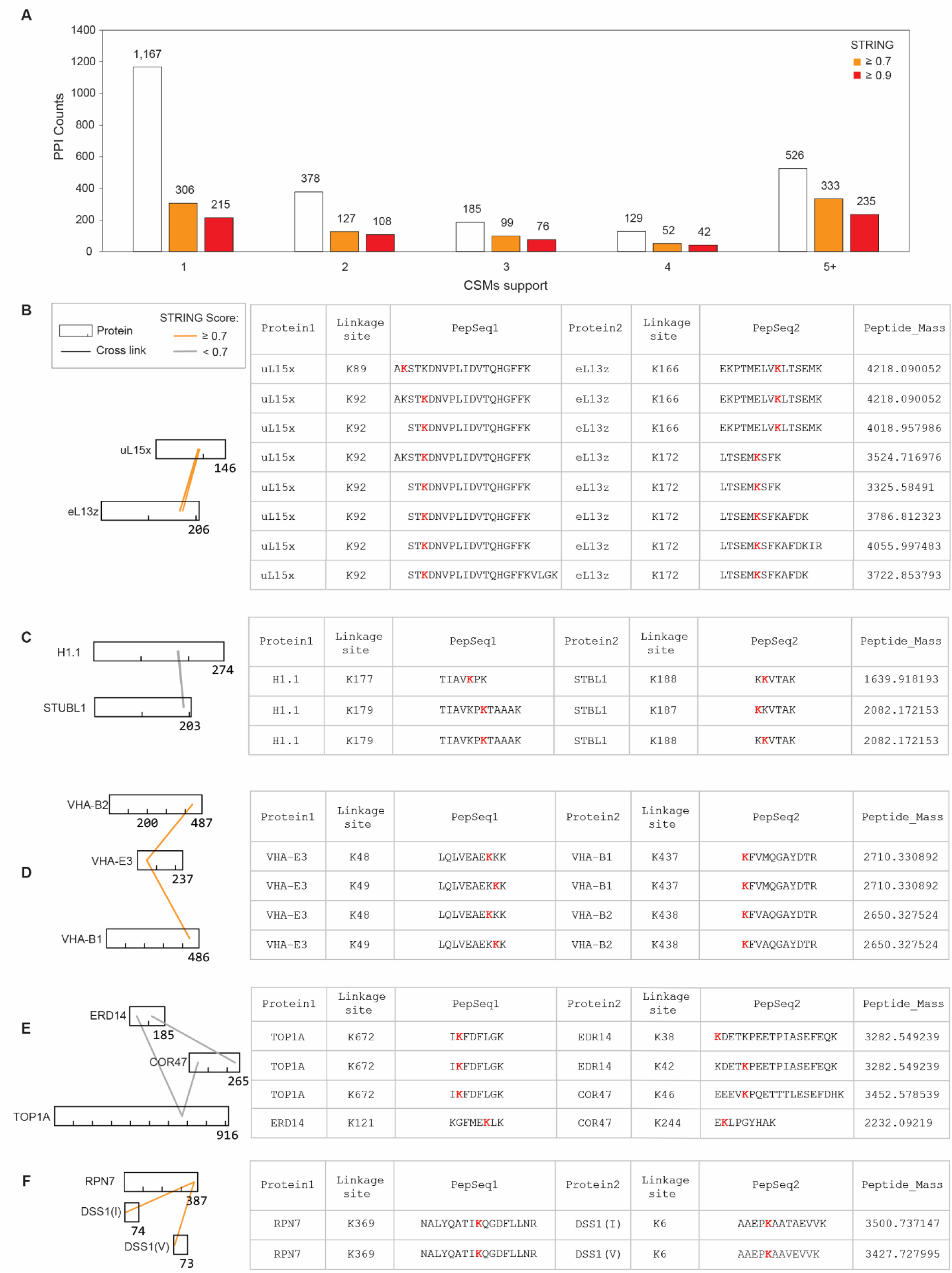
CSM support for PPI identifications and alignment of observed cross links with site specific and homologous protein regions. Evidence of identification consistency is shown by redundant mapping to identical sites or conserved regions. Precursor mass, peptide sequence, and cross-linked sites are shown. (**A**) Distribution of CSM support for PPI identifications. A total of 1,167 heteromultimeric interactions (49%) were supported by a single cross-linked spectral match (CSM), while 51% were supported by multiple CSMs. Only uniquely identified CSMs were included in the analysis, with redundant matches removed. A CSM was defined as distinct based on differences in peptide backbone sequence, cross-linking site, charge state, or variable modifications (e.g., oxidation). STRING scores are provided for comparison. The x-axis represents unique CSM counts, and the y-axis represents PPI counts. (**B**) Ribosomal subunit cross-links between uL15x K92 and eL13z-K166 or K172, showing variable miscleavages. (**C**) Interaction between Histone 1.1 K177 or K179 and SUMO targeted ubiquitin E3 ligase 1, STUBL1, at K187 or 188. (**D**) Vacuolar proton ATPase subunit VHA-E3-K49 linked to homologous sites VHA-B1 K437 and VHA-B2 K438. (**E**) DNA topoisomerase I alpha, TOP1A, K672 linked to dehydrin homologs ERD14 at K38 or K42 and COR47 at K46. (**F**) 26S proteasome subunit RPN7-K369 linked to the same sites in homologs DSS1(I) K6 and DSS1(V) K6.

**Supplementary Fig. S5.**
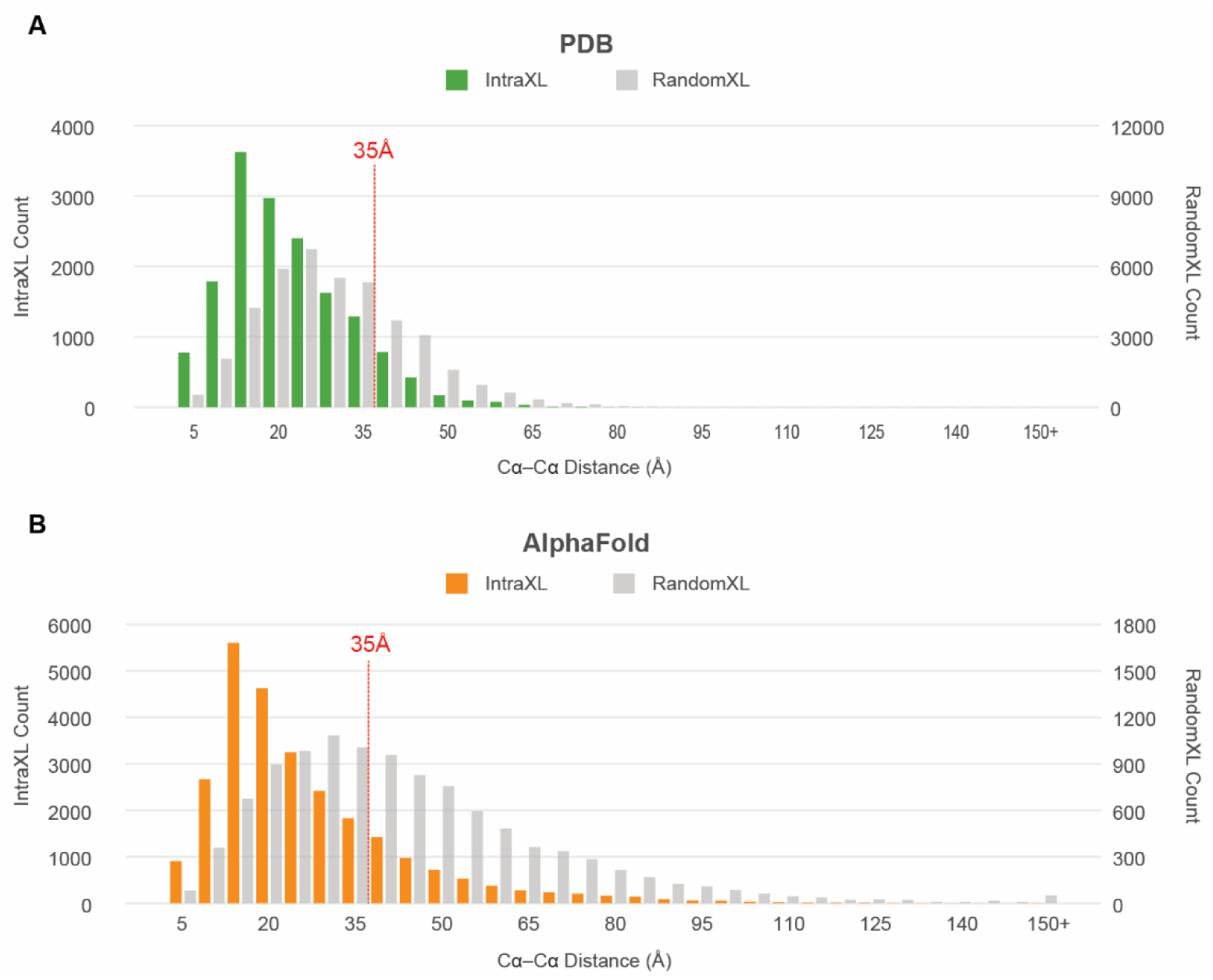
Validation of cross-link distance constraints using PDB and AlphaFold structures. (**A**–**B**) Distributions of minimum Euclidean distances (Å, Cα–Cα) for identified cross-linked residue pairs compared with randomly generated residue pairs mapped onto Protein Data Bank (PDB) structures (A) and AlphaFold models (B) from *Arabidopsis*. Notably, the mean and maximum distances observed in AlphaFold structures are greater than those in PDB structures, likely reflecting the fact that PDB entries often represent partial protein fragments rather than full-length proteins.

**Supplementary Fig. S6.**
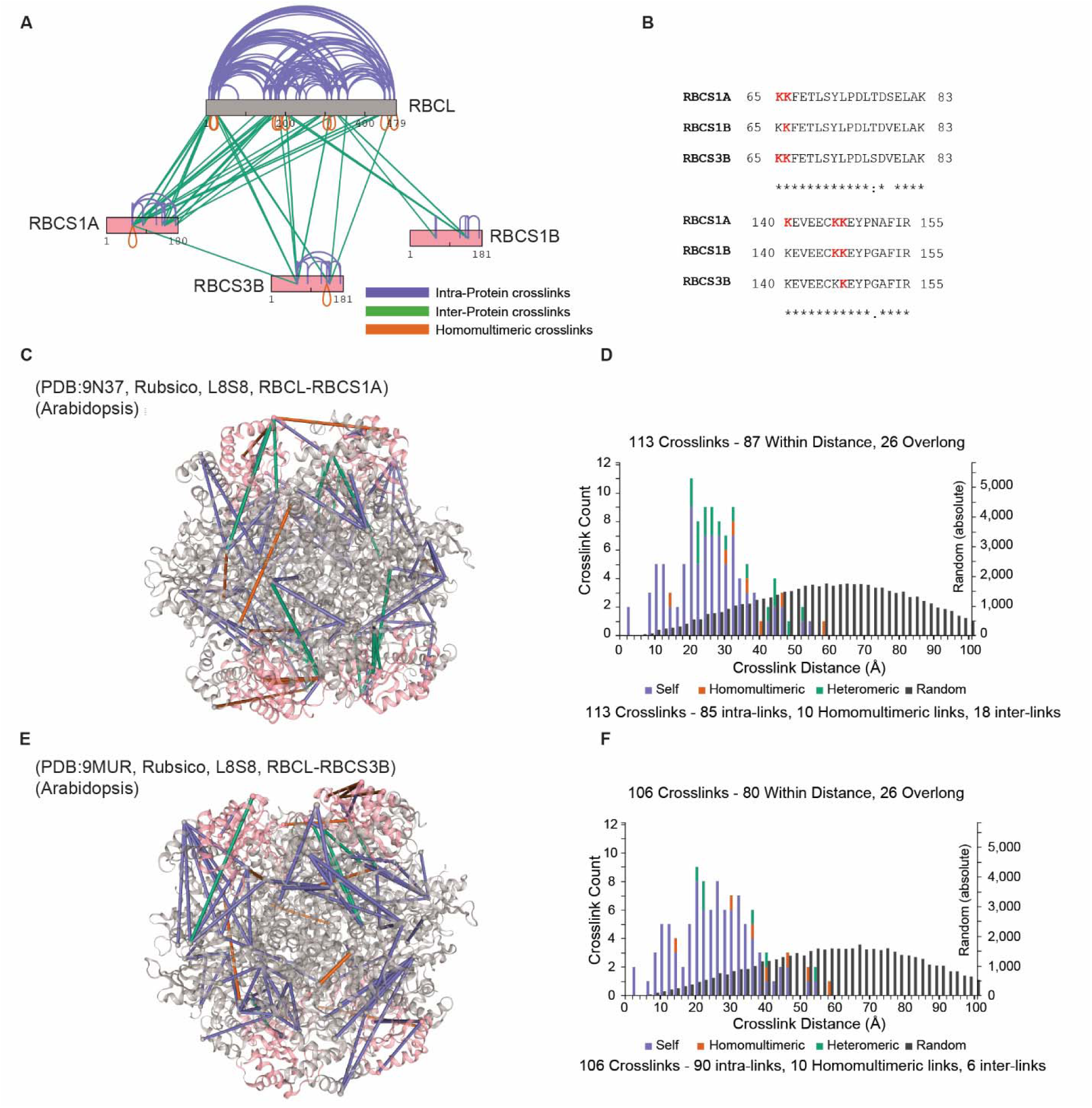
Mapping of Rubisco cross-links on PDB structures of Arabidopsis Rubisco. Cross-links are color-coded by type: purple for self (intra)links, green for heteromeric (inter)links, and orange for homomultimeric links. (**A**) Schematic of Rubisco Cross-linking: Diagram illustrating intra-protein cross-links within and inter-protein cross-links between the large subunit (RBCL, grey) and three small subunit isoforms (RBCS1A, 1B, and 3B, pink). RBCS1A is the dominant isoform and therefore yielded the highest number of identified cross-links with RBCL. (**B**) Sequence Alignment and Isoform-Specific Cross-links: Multiple sequence alignment of RBCS1A, 1B, and 3B highlighting regions (residues 65–83 and 140–155) containing identified cross-linked peptides with isoform-specific sequence variations. Lysine residues (K), representing cross-linking sites, are highlighted in red. Only peptides with sequence differences are displayed to demonstrate the specificity of isoform assignments. Conservation is indicated by asterisks (*, identical), colons (:, strongly similar), and periods (., weakly similar). (**C**–**D**) RBCL–RBCS1A Mapping: Cross-links between RBCL and RBCS1A mapped onto the Arabidopsis L_8_S_8_ Rubisco structure (PDB: 9N37), with the accompanying histogram showing the Cα-Cα distance distribution. RBCL in the 9N37 structure lacks C-terminal region (residues 459-479), reducing cross-link mapping coverage. (**E**–**F**) RBCL–RBCS3B Mapping: Cross-links between RBCL and RBCS3B mapped onto the Arabidopsis L_8_S_8_ Rubisco structure (PDB: 9MUR), with the accompanying histogram showing the Cα-Cα distance distribution. RBCL in the 9MUR structure lacks C-terminal region (residue 465-479), reducing cross-link mapping coverage.

**Supplementary Fig. S7.**
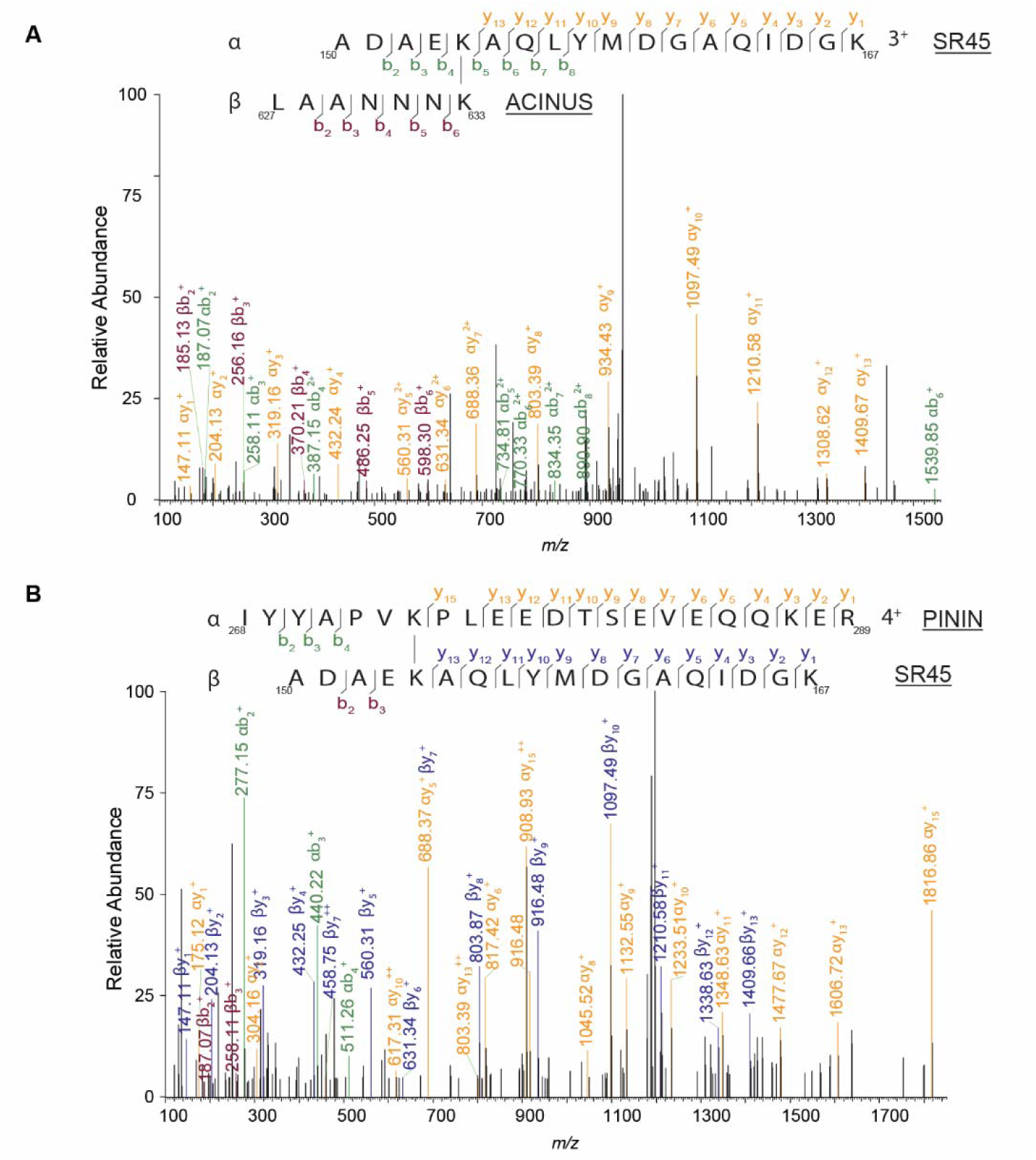
High-confidence identifications of cross-linked peptides between ACINUS–SR45 and PININ–SR45. b-ions are shown in green and y-ions in yellow. A series of fragment ions, including those from α- and β-peptides, as well as cross-linker-containing ions, were detected, supporting the reliability of the cross-link identifications. (**A**) Spectrum of the cross-linked peptide between ACINUS-K633 and SR45-K154. (**B**) Spectrum of the cross-linked peptide between PININ-K274 and SR45-K154.

**Supplementary Fig. S8.**
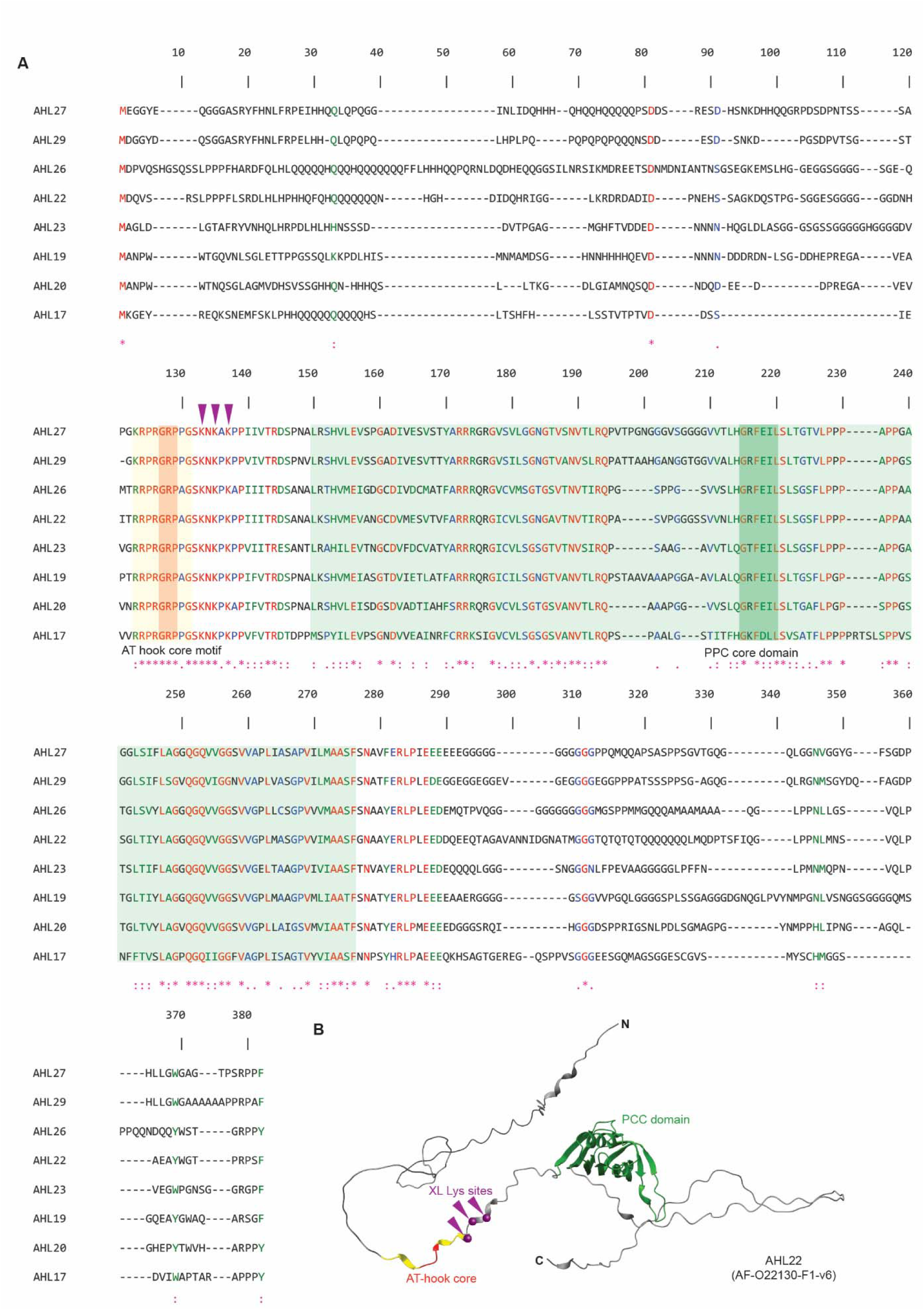
Sequence Alignment of the AHL Family and AlphaFold Structural Modeling of Crosslinked Lysine Sites. (**A**) Multiple Sequence Alignment of AHL Family Members. The alignment was performed using ClustalW. The AT-hook core motif and the Plant and Prokaryote Conserved (PPC) domain (including the core domain) are highlighted in yellow and green, respectively. Within these regions, the core GRP is indicated in purple, and the core GRFEIL is shown in dark green. Red arrows denote the positions of crosslinked lysine residues. (**B**) AlphaFold Structural Prediction of AHL22. The predicted structure highlights the AT-hook core in yellow and the PPC structured domain in green. Three specific crosslinked lysine sites are indicated by arrows.

**Supplementary Fig. S9.**
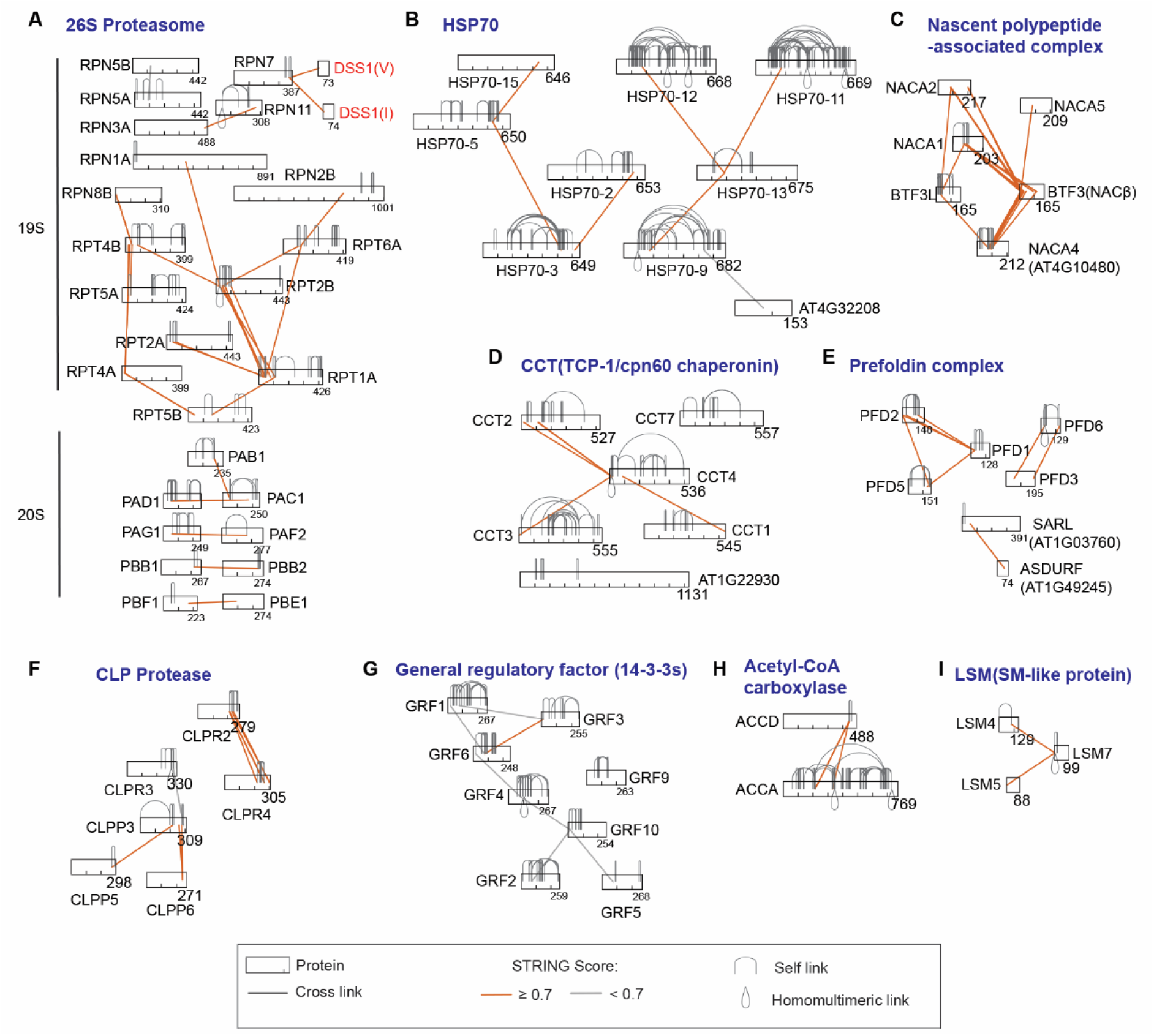
Topological mapping of multiple protein complexes by XL-MS. Proteins are represented as rectangles scaled according to their respective lengths. Cross-links are color-coded based on STRING confidence score thresholds, highlighting both known and novel interactions. (**A**) Topological mapping of the 26S proteasome complex, comprising the 19S and 20S subunits, showing connectivity among subunits. Interaction interfaces between RPN7 and DSS1(I) and DSS1(V) are highlighted. DSS1 proteins are particularly challenging to resolve by cryo-EM because of their small size, intrinsic disorder, and dynamic interaction properties. (**B**–**I**) Topological mapping of multiple protein complexes, including several chaperone-related complexes such as Heat Shock Protein 70 (HSP70), the Nascent Polypeptide-Associated Complex (NAC), CCT (TCP-1/cpn60 chaperonin), Prefoldin (PFD), CLP protease, and General Regulatory Factors (14-3-3 proteins). Additional complexes include Acetyl-CoA Carboxylase (ACC) and the Sm-like small nuclear ribonucleoprotein (LSM) complexes.

**Supplementary Fig. S10.**
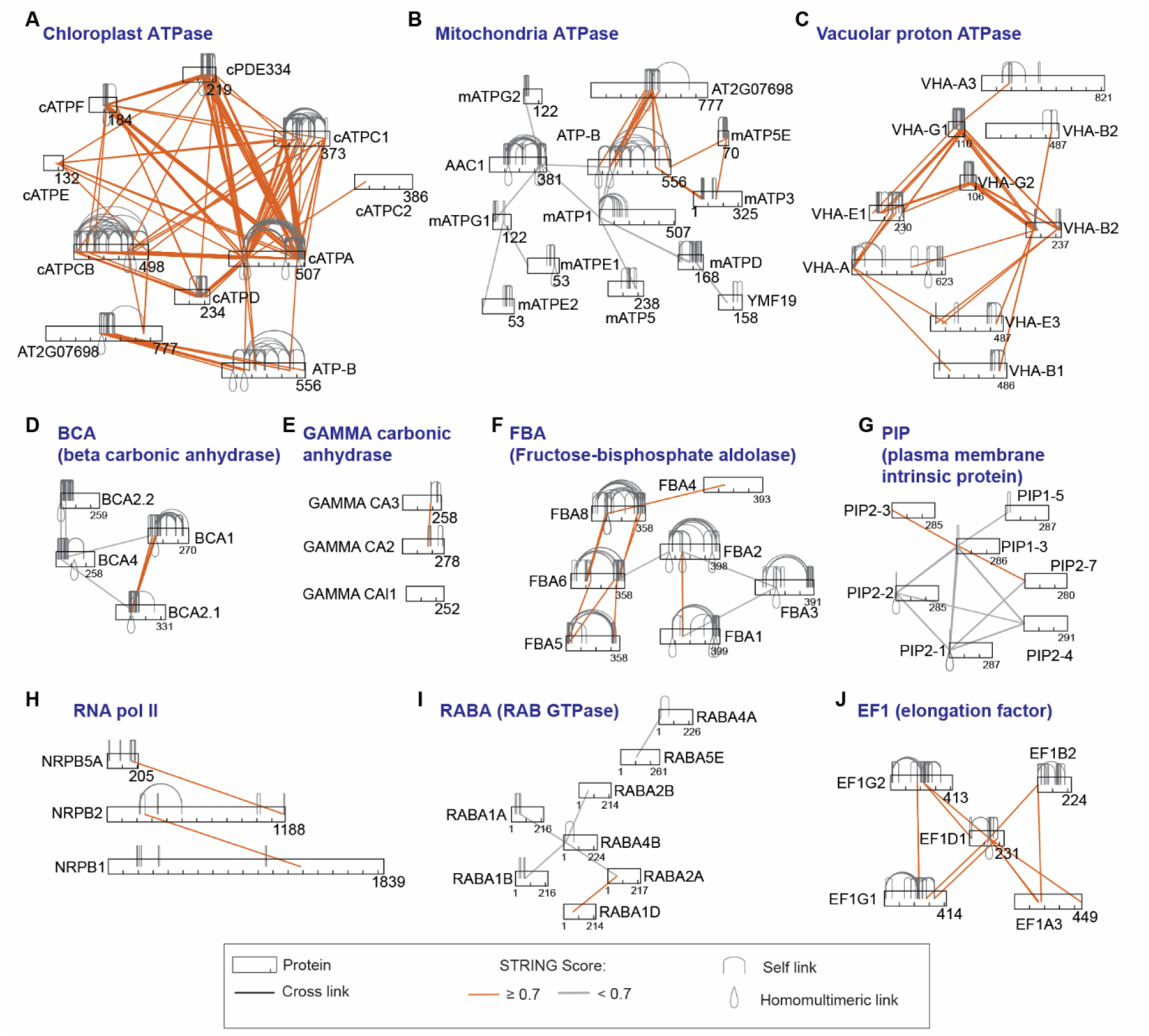
Topological mapping of multiple protein complexes by XL-MS. (**A**-**C**) Topological mapping of ATPase complexes from chloroplasts, mitochondria, and vacuolar proton ATPase. AT2G07698, encoding the ATPase F1 complex α-subunit, and ATP-B (AT5G08670), both nuclear-encoded proteins, formed cross-links with chloroplast-encoded ATPase complex subunits as well as mitochondria-encoded ATPase subunits, suggesting dual localization and assembly in both organelles. (**D**-**E**) Topological connections of β- and γ-carbonic anhydrases. (**F**-**J**) Topological mapping of multiple protein complexes, including fructose-bisphosphate aldolase, plasma membrane intrinsic protein complexes, RNA polymerase II, RAB GTPase, and elongation factor complexes.

**Supplementary Fig. S11.**
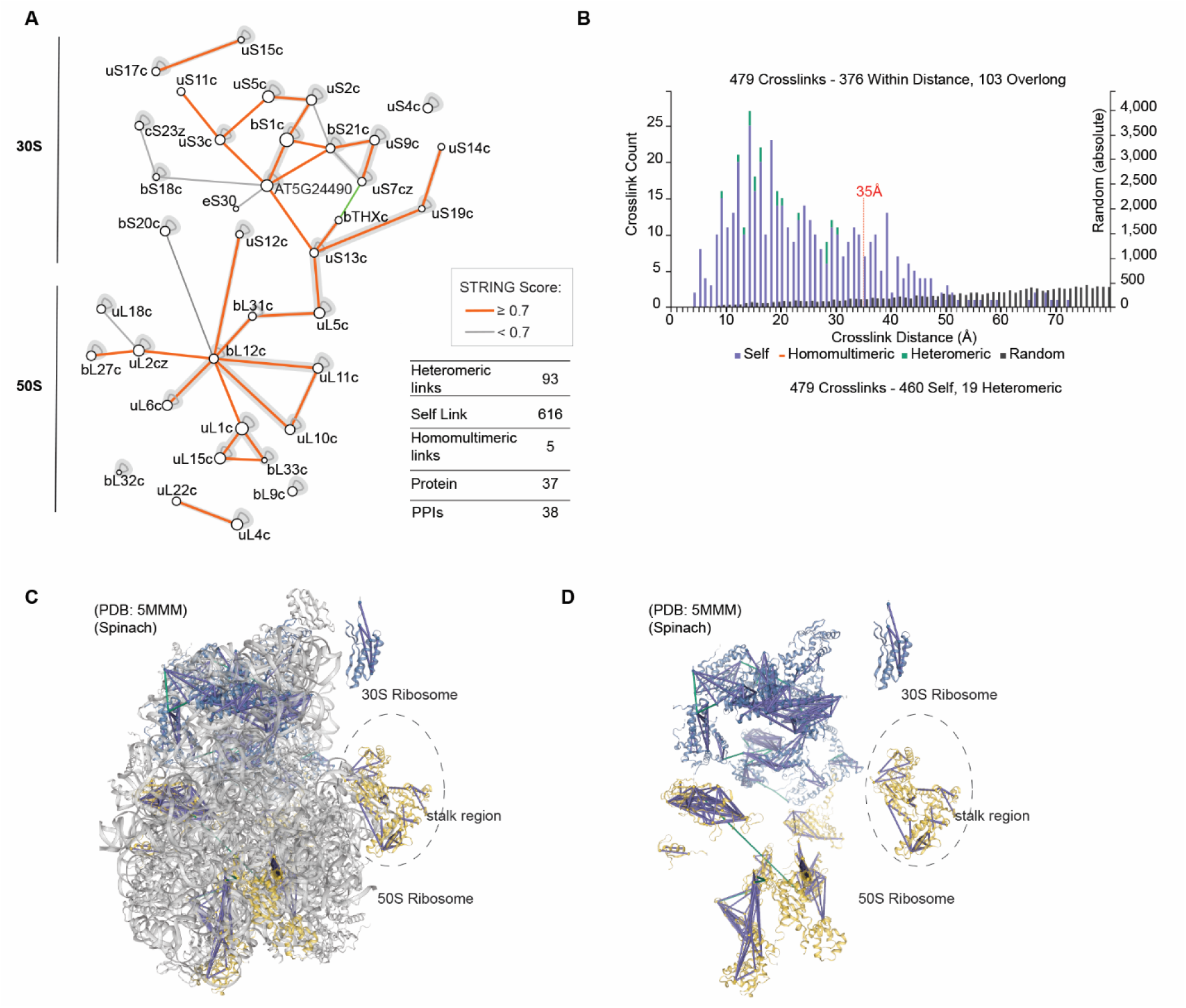
Mapping of cross-links in the chloroplast 70S ribosome complex onto the PDB structure of spinach 70S ribosomes. (**A**) Topological mapping of cross-links within the 70S ribosome complex. (**B**–**D**) A total of 479 cross-links were mapped onto the spinach cryo-EM structure (PDB: 5MMM), which lacks bL12c in the stalk region. The structure contains both ribosomal subunits and associated ribosomal RNAs. In (D), ribosomal RNAs were removed to facilitate clearer visualization of the cross-links. The accompanying histogram in (B) shows the distribution of Cα–Cα distances for the mapped cross-links.

**Supplementary Figure S12.**
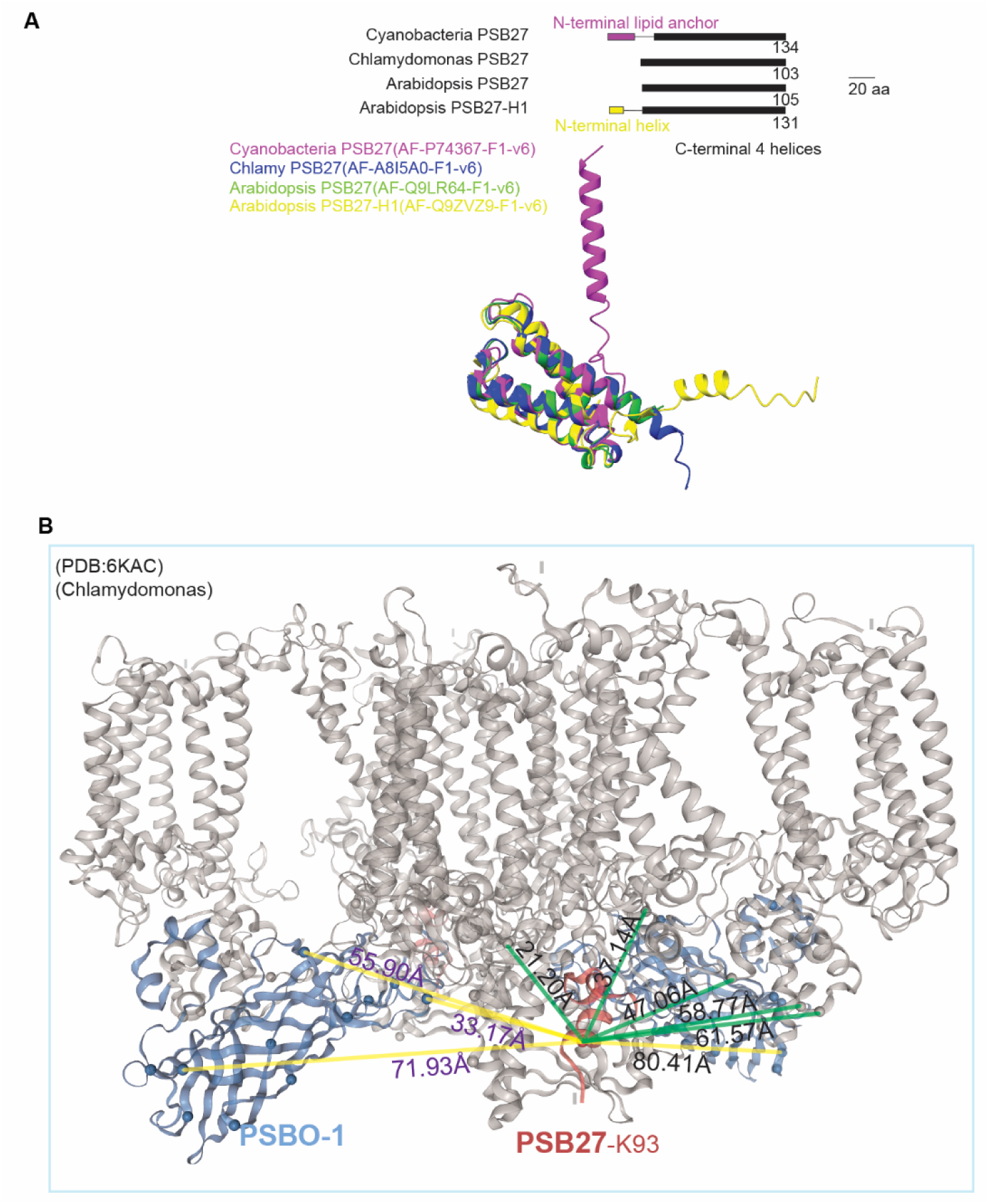
Mapping of PSB27 cross-links onto PDB structure 6KAC from Chlamydomonas. (**A**) Superimposed AlphaFold structures of PSB27 homologs reveal unique structural features of Arabidopsis PSB27. Although Arabidopsis PSB27 retains the conserved C-terminal four-helix bundle, it lacks the lipid anchor and displays divergence in both sequence and surface charge distribution. (**B**) Cross-links were mapped onto the Chlamydomonas PSII structure (PDB: 6KAC), which contains residues 91–114 of PSB27. PSB27 is shown in purple, PSII core subunits in gray, and the PSBO-1 protein of the oxygen-evolving complex (OEC) subunits in blue. Yellow lines indicate cross-links between PSB27 and PSBO-1, whereas green lines indicate cross-links between PSB27 and PSII core subunits (PsbA, PsbB, PsbC, and PsbD). The observed distance constraints suggest a dynamic interaction between PSB27 and the PSII complex.

**Supplementary Fig. S13.**
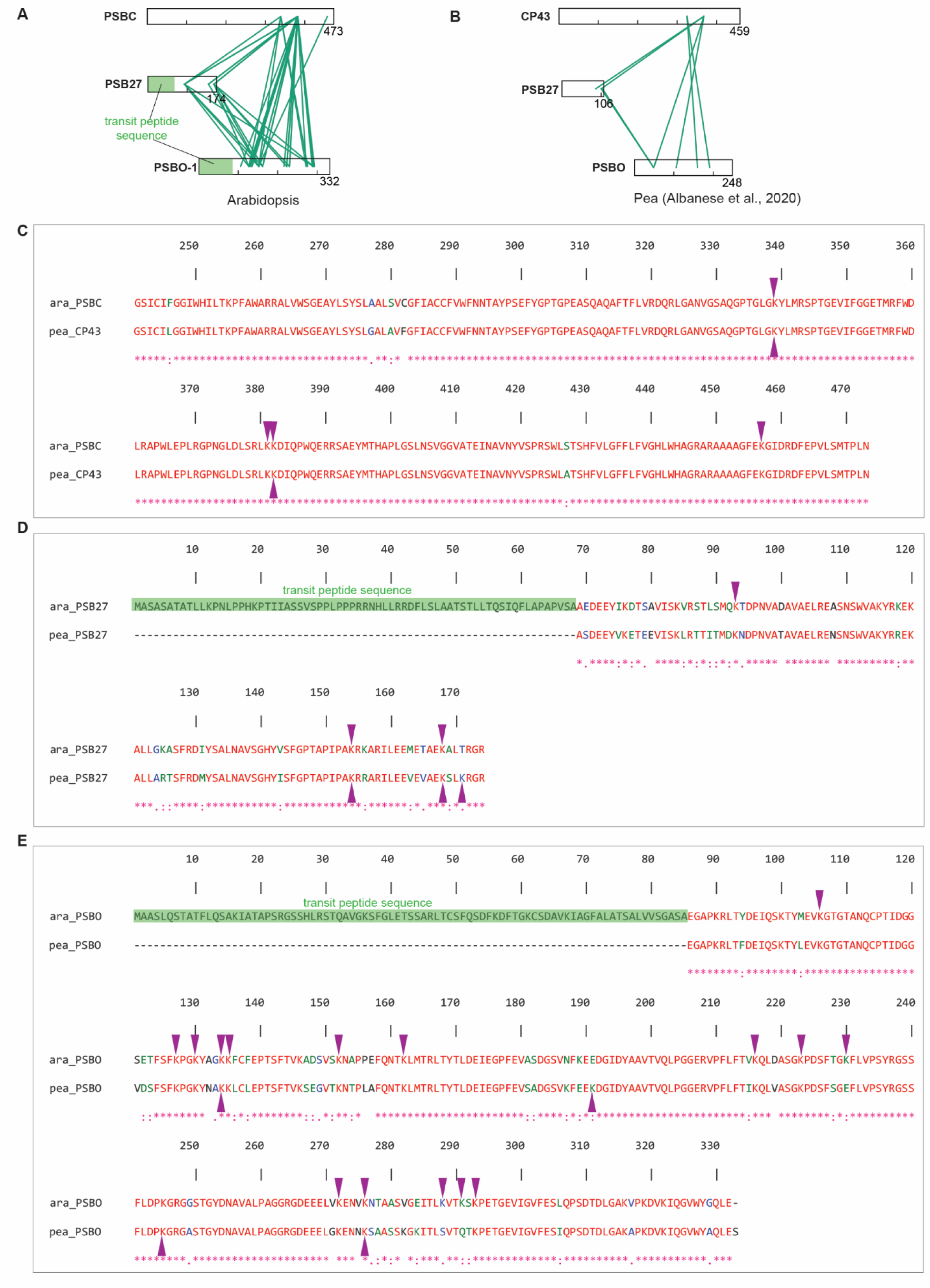
Comparison of cross-linked sites among PSB27, PSBC, and PSBO between Arabidopsis and pea. (**A**–**B**) Detected cross-links between PSB27 and PSBO, and between PSB27 and PSBC, in Arabidopsis and pea (Albanese, 2020). (**C**–**D**) Comparison of the detected cross-linked sites (indicated by arrows) between the two species, revealing conserved interaction sites. Additional cross-linked sites were identified in the present study.

**Supplementary Fig. S14.**
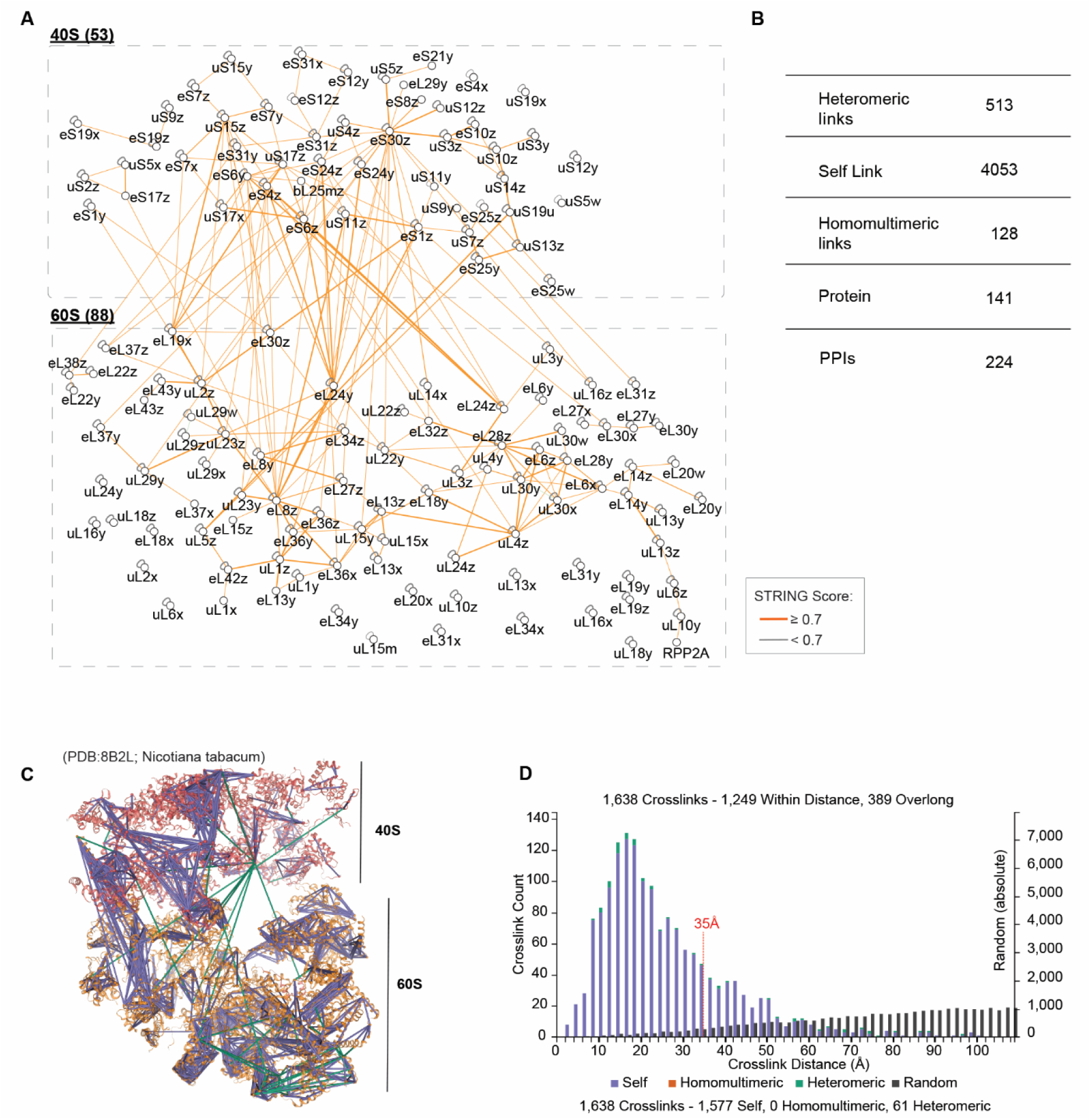
Topology of the 80S ribosomal complex. (**A**) Topological arrangement of the 40S and 60S ribosomal subunits based on their connectivity. Links between proteins are colored according to the confidence thresholds of the STRING score. (**B**) Summary of cross-links and protein-protein interaction (PPI) information from (A). (**C**) 80S ribosome cross-links mapped onto tobacco 80S ribosome (PDB: 8B2L). The large ribosomal subunit is colored orange and the small ribosomal subunit is colored red. Links are color-coded according to link type: purple for self-links (intralinks), green for heteromeric links (interlinks), and orange for homomultimeric links. Ribosomal RNAs were removed to facilitate clearer visualization of the cross-links (**D**) Distribution of cross-link distances from the integrated cross-link model in (C).

**Supplementary Fig. S15.**
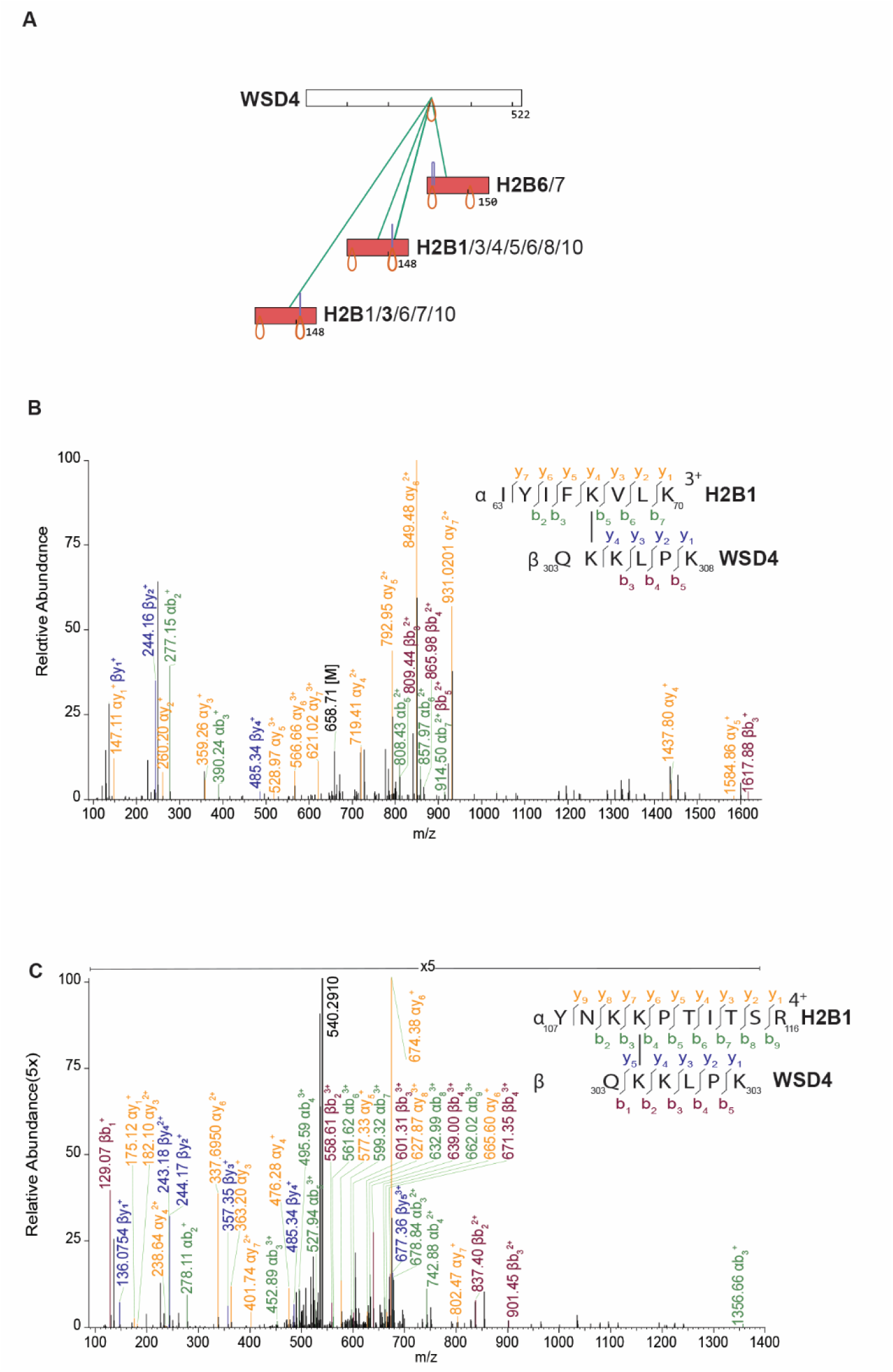
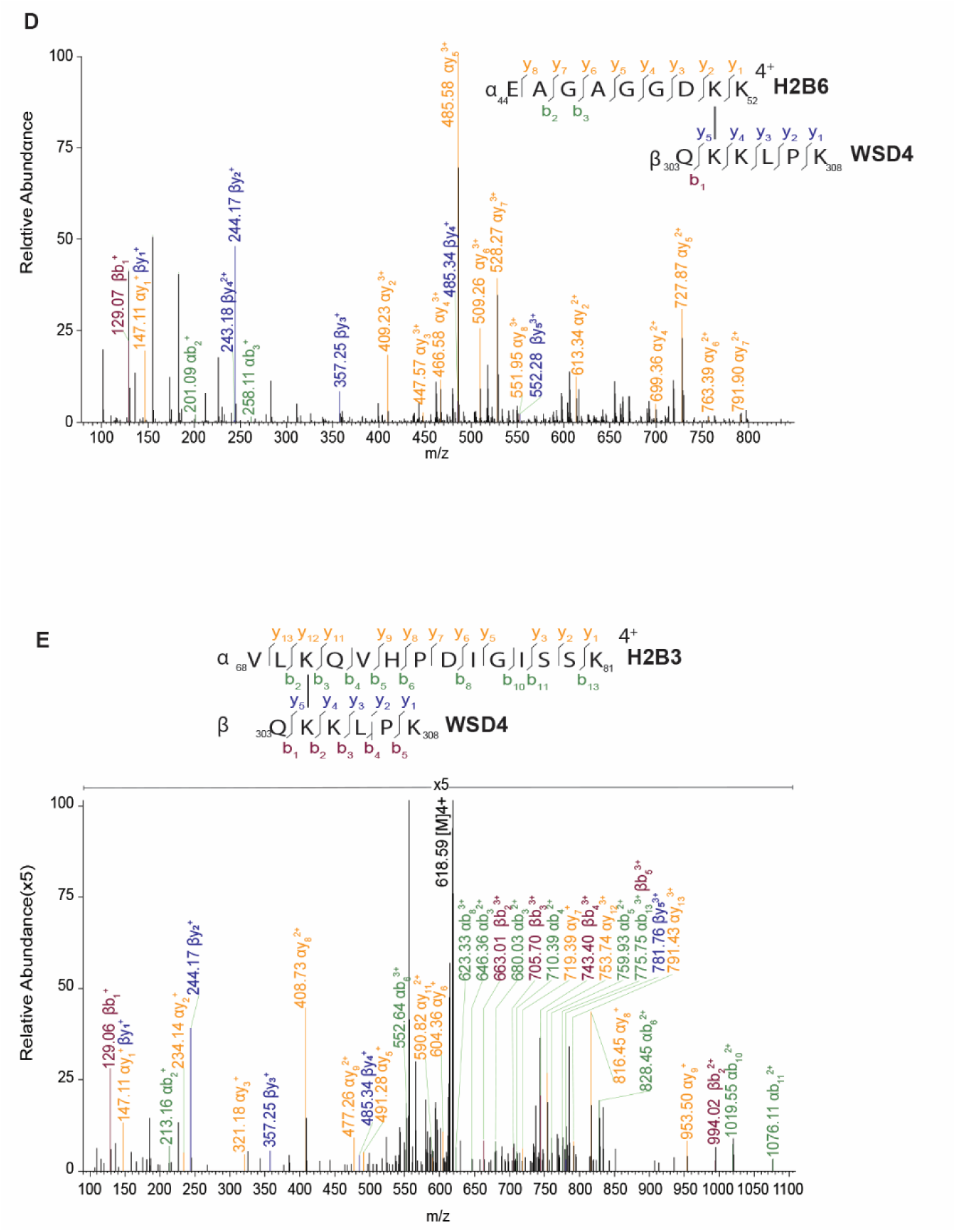
XL-MS identifies WSD4 as a novel histone interactor associated with H2B family members. (**A**) Cross-links were detected between WSD4, an O-acyltransferase, and H2B family members. Owing to the high sequence homology among H2B family proteins, several identified peptides mapped to shared H2B sequences. (**B**–**E**) Representative cross-link spectra identified by pLink3.2 demonstrating interactions between WSD4 and H2B family members. Multiple series of fragment ions, including α- and β-peptide ions as well as cross-linker-containing fragment ions, were detected, supporting the confidence and reliability of the cross-link identifications.

**Supplementary Fig. S16.**
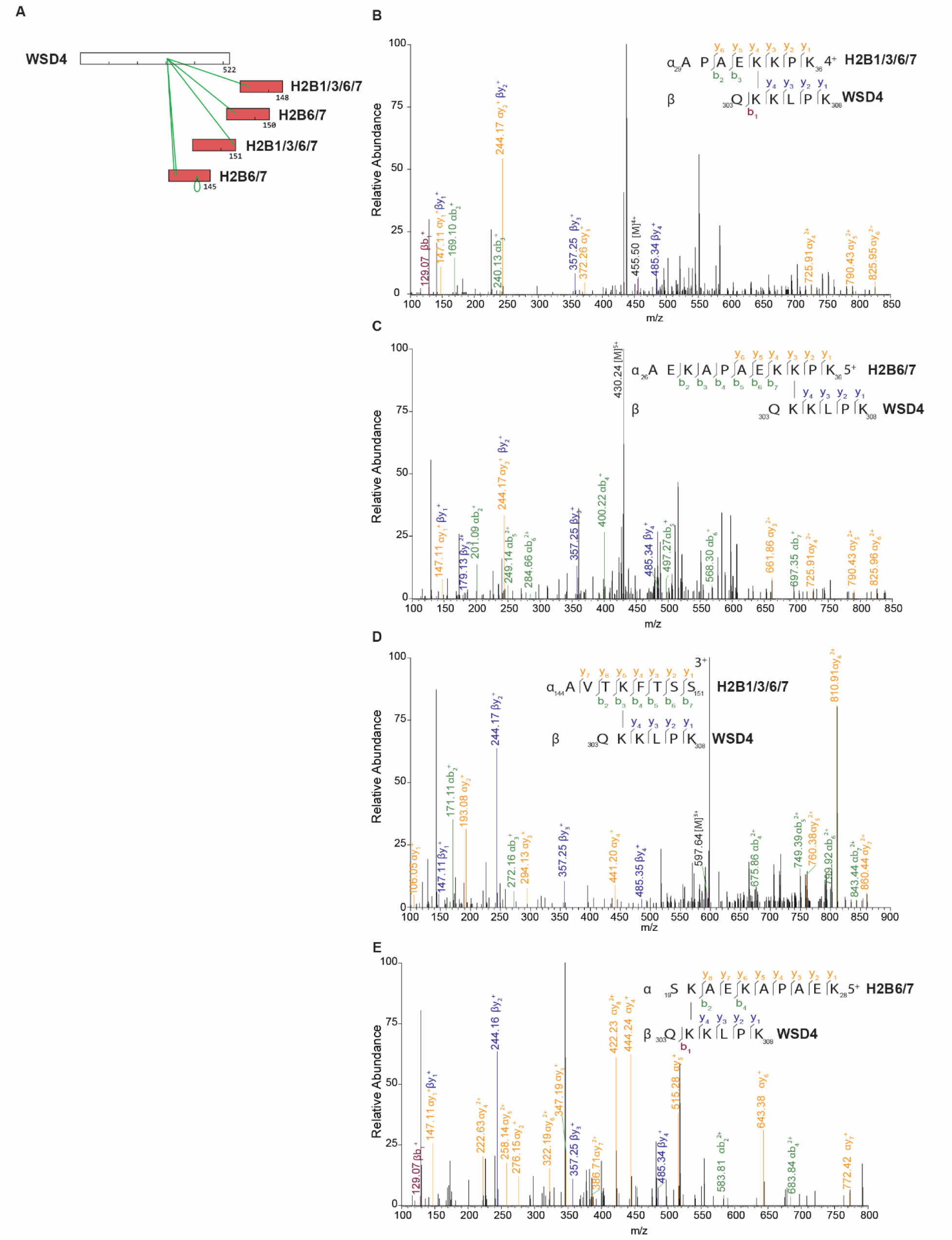
Supplementary Fig. S16. XL-MS identifies WSD4 as a novel histone interactor associated with H2B family members using pLink2. (**A-E**) Cross-linked peptides identified by pLink2.3.11 at a 5% FDR at the CSM level support the PPI between H2B family members and WSD4. These peptides were not identified by pLink3.2.0 at a 1% peptide-level FDR.

**Supplementary Fig. S17:**
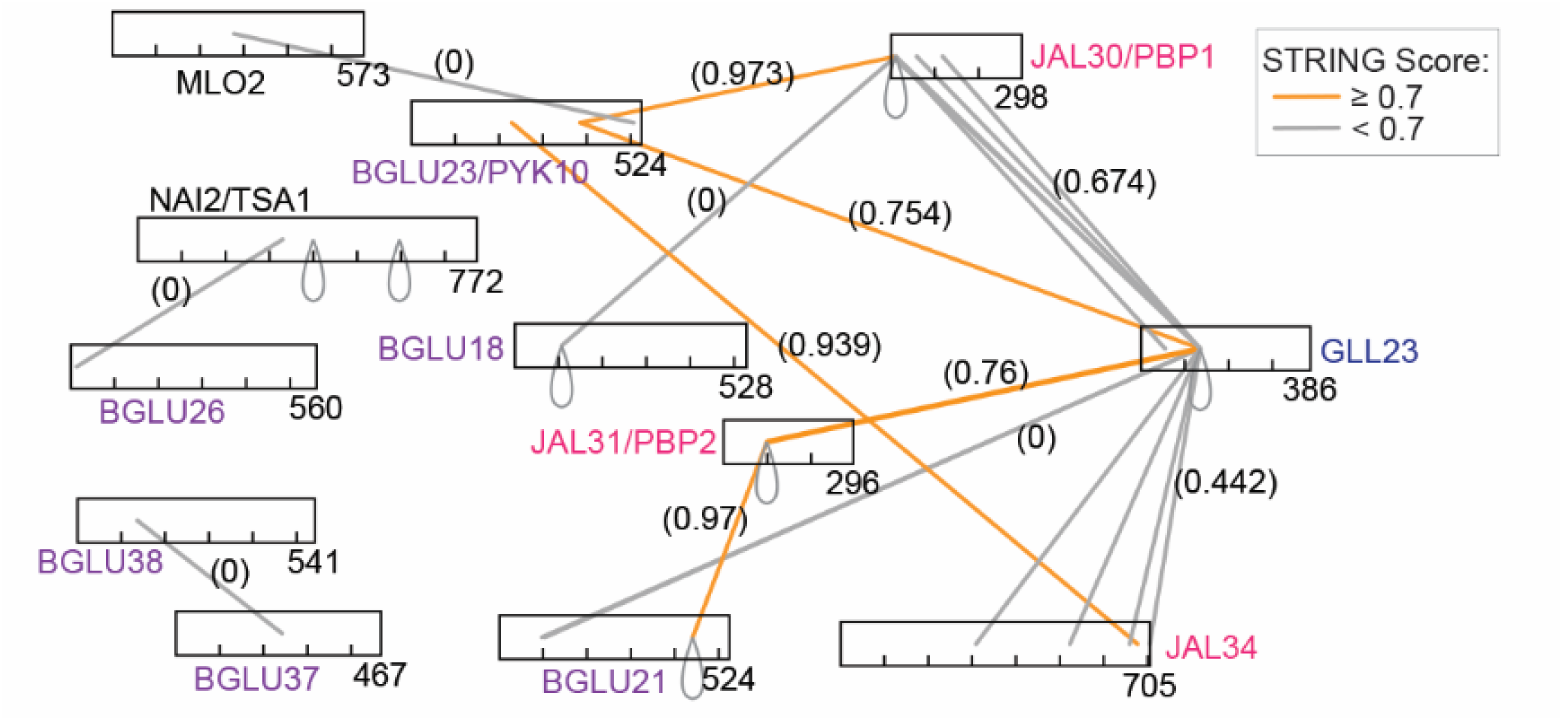
Interaction network of a specialized defense system detected by XL-MS. XL-MS analysis revealed an extensive interaction network spanning multiple protein families involved in a specialized defense system. The network highlights extensive connectivity among BGLU family members, major constituents of ER bodies that encode β-glucosidases, together with jacalin-related lectin JAL/PBP proteins, which may function as molecular chaperones that facilitate proper BGLU polymerization, and GLL lipase-like proteins proteins. Additional interactions were detected with NAI2, a master regulator of ER body formation, and MLO2, a homolog of the barley Mildew Resistance Locus O protein. Detection of both heteromeric and homomultimeric crosslinks suggests the formation of higher-order protein assemblies that may enable rapid activation of defense components upon cellular wounding. Although GLL23 proteins are normally localized in the cytoplasm, cellular wounding promotes their interaction with ER body proteins, including BGLUs and JAL/PBP proteins. Lines are color-coded according to STRING confidence scores (orange, ≥0.7; grey, <0.7), with individual scores indicated in parentheses.

**Supplementary Fig. S18.**
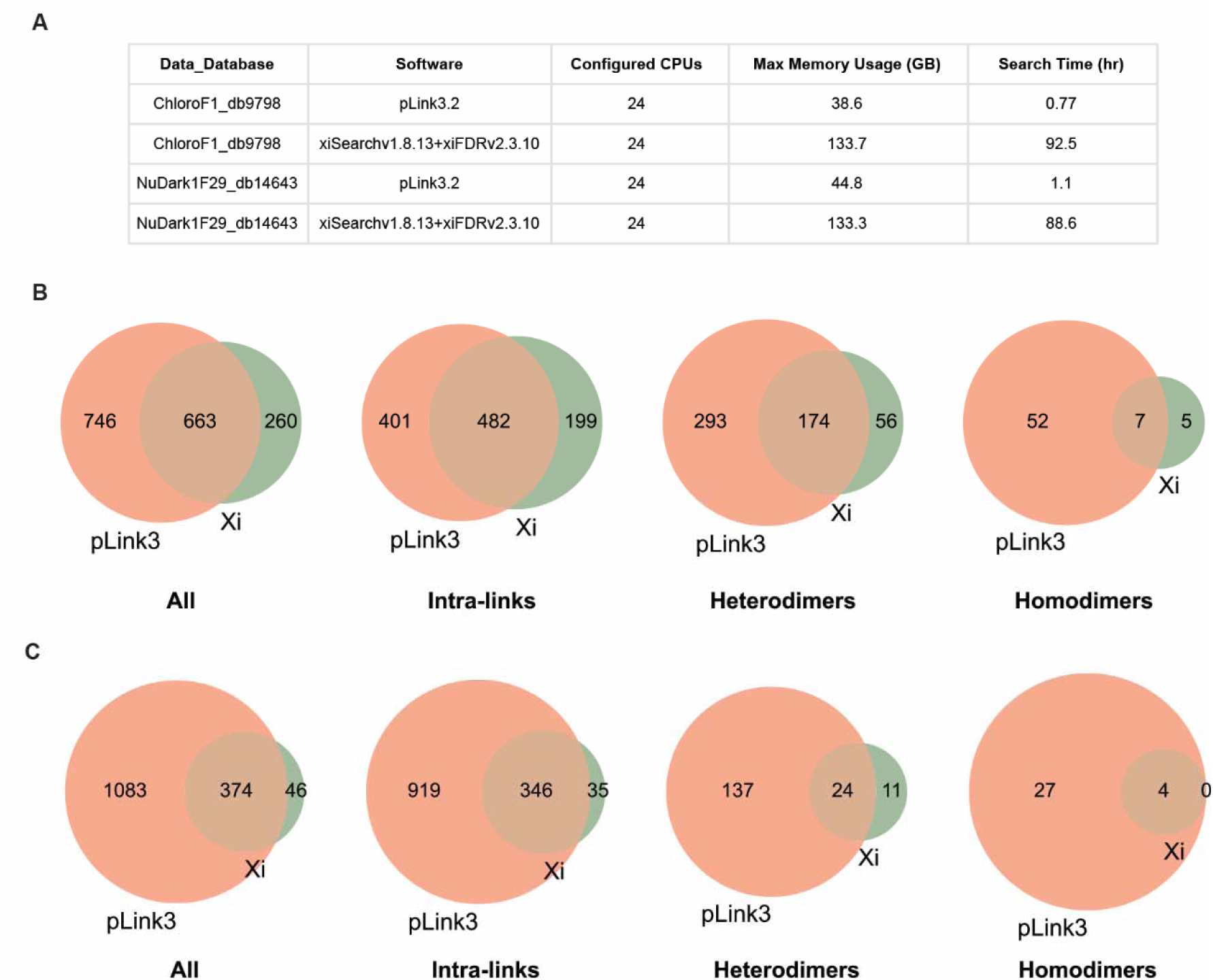
Comparison of data searches performed using pLink3 and xiSEARCH. (**A**) Comparison of memory usage and search time between pLink3 and xiSEARCH using a single raw XL-MS data file. As xiSEARCH required longer processing times under our computational settings, whole-dataset comparisons were not performed. Instead, comparisons were conducted using two representative fractionation datasets: one chloroplast fraction and one nucleus fraction. (**B**–**C**) Comparison of cross-linked peptide identifications obtained by pLink3 and xiSEARCH from single raw XL-MS datasets derived from chloroplast (B) and nucleus (C) fractions. A large proportion of cross-linked peptides were identified by both search engines, while each also detected unique cross-linked peptides. Overall, pLink3.2 identified a greater number of XL peptides.

